# Diverse Ligands for Mycobacterial CYP124 Identified from Plant and Marine Compounds

**DOI:** 10.64898/2026.01.23.701388

**Authors:** Leonid Kaluzhskiy, Evgeniy Yablokov, Tatsiana Varaksa, Anna Grudo, Alena Karputs, Irina Grabovec, Yuri Mezentsev, Elena Zelepuga, Oksana Gnedenko, Dmitriy Tolkanov, Timofey Malyarenko, Sophia Kolesnikova, Anastasia Kozhushnaya, Elena Vasileva, Natalia Mishchenko, Alexandra Silchenko, Sergey Avilov, Tatyana Rutckova, Darya Tarbeeva, Vladimir Kalinin, Alla Kicha, Emma Kozlovskaya, Sergey Fedoreyev, Natalia Ivanchina, Pavel Dmitrenok, Andrei Gilep, Alexis Ivanov

## Abstract

Tuberculosis persists as a major global health threat, significantly exacerbated by the rise of drug-resistant strains. Cytochrome P450 of 124 family CYP124 from Mycobacterium tuberculosis (CYP124), implicated in host sterol metabolism and bacterial virulence, represents an emerging and promising therapeutic target. While its precise physiological role was previously debated, CYP124’s confirmed ability to metabolize immunomodulatory host sterols underscores its pharmacological relevance. Utilizing surface plasmon resonance binding assays and UV-Vis spectral titration screening, we identified nine novel non-azole ligands for CYP124 from a library of 32 plant-derived and marine natural compounds. Among these hits, (25S)-5α-cholestane-3β,4β,6α,7α,8,15β,16β,26-octaol (termed 15β-octaol) and henricioside H_2_ (HD-4) induced characteristic difference spectra and formed long-lived inhibitory complexes with CYP124, exhibiting dissociation half-lives of 181 min and 65 min, respectively. However, their inhibitory potency was moderate, with IC_50_ values of approximately 86 μM for 15β-octaol and exceeding 100 μM for HD-4. Complementary *in silico* molecular docking and analysis identified key conserved hydrophobic residues within the CYP124 active site crucial for ligand binding, suggesting a shared pharmacophore. Furthermore, structural similarity analysis revealed that 37 human endogenous metabolites, including known immunoregulatory sterols, bear resemblance to the identified CYP124 ligands. This finding points towards a potential sterol-mediated interplay at the host-pathogen interface. Collectively, these results provide a foundation for the future development of mechanism-based CYP124 inhibitors as therapeutics against multidrug-resistant tuberculosis.

## 1. INTRODUCTION

Tuberculosis (TB), an infection caused by *Mycobacterium tuberculosis* (Mtb), currently represents one of the pressing problems in medicine. According to WHO data in 2023, 10.8 million new cases of TB were registered, while 1.25 million people died from this infection. Thus, tuberculosis has become the leading infectious cause of mortality. Furthermore, the prescription of adequate drug therapy remains a major challenge. Therapy for multidrug-resistant TB (MDR-TB) presents particular complexity. According to WHO data, only about 40% of patients with drug-resistant tuberculosis received treatment in 2023[1]. Consequently, the search for potential therapeutic agents active against Mtb represents a highly relevant biomedical task. Simultaneously, the search for new Mtb target proteins for pharmacological intervention is pertinent [2,3].

Mtb cytochrome P450s (enzymes of the heme-thiolate monooxygenase superfamily) are promising targets for TB therapy[4,5]. Classical inhibitors of these enzymes are compounds of the azole class [6], for which antibacterial activity, including against multidrug-resistant strains of Mtb, has been demonstrated[7,8]. Concurrently, promising non-azole inhibitors of Mtb cytochrome P450s are being investigated[5,9–14]. Since Mtb possesses multiple cytochrome P450s, they are ranked according to their suitability as drug targets. According to Ortiz de Montellano [4], CYP121, CYP125, and CYP142 are potential drug targets; however, the author emphasises that CYP124 is not a clear target for the development of anti-tuberculosis drugs[4] due to the lack of sufficient data. Hudson et al.[5] declared CYP121, CYP125, and CYP128 as the most attractive anti-TB targets, while CYP124 and CYP142 were mentioned as secondary targets. Ortega Ugalde [15] declared CYP121A1, CYP125A1, CYP139A1, CYP142A1, and CYP143A1 as anti-TB targets. It is known that Mtb possesses a powerful sterol catabolism system crucial for its virulence[16]. This system, which includes the sterol-oxidising CYP124, is currently considered a promising target for pharmacological intervention[17,18]. The key hypothesis is that CYP124, participating in ensuring Mtb virulence (although not being vitally essential *in vitro*), plays an important role in the interaction of the pathogen with host biochemistry[19]. Specifically, the enzyme catalyses ω-hydroxylation of methyl-branched lipids (e.g., phytanic acid), which can modulate the immune response or bacterial cell wall formation *in vivo*[20], and also transforms human immunomodulatory sterols[21–23]. Thus, despite the controversial status of CYP124 as a primary target, its role in the metabolism of host immunoregulatory sterols makes the enzyme a promising target for suppressing Mtb virulence.

Based on this hypothesis, the search for new potential ligands for the CYP124 active site represents a promising direction. Within this task, two complementary approaches can be distinguished: screening natural compounds as a traditional source of therapeutic agents and searching for new potential ligands for the CYP124 active site among human endogenous metabolites, based on its physiological function. The latter can be implemented using computational screening methods, including analysis of chemical similarity to known substrates and ligands.

Natural compounds and their analogues have historically served as the basis for therapeutic agents [24], a resource documented in numerous dedicated databases[25,26]. Plants are the most popular source of natural compounds[27]. One well-studied example is *Maackia amurensis*, considered by many authors as a source of potential therapeutic compounds[28–30]. In the work of Noviany Hasan et al., the inhibitory effect on Mtb of medicarpin[31], an isoflavonoid from some plants, including *M. amurensis*[32], was studied. Marine animals and plants also attract significant interest as sources of compounds possessing diverse biological activity. For example, compounds isolated from seagrass of the species*Zostera marina,* exhibit a wide spectrum of activities, including antioxidant, antifungal, and antiviral[33], and extracts of its microbiome have potential for the search for new antibiotics[34]. In addition to marine plants, representatives of the world ocean fauna are currently considered as promising sources of natural compounds, for example, representatives of the phylum Echinodermata[35–39]. In general, compounds isolated from Echinodermata can be considered as potential agents against Mtb[40]. Thus, some holothurins and saponins isolated from sea cucumbers have shown activity against Mtb, including against drug-resistant strains [41–44]. Besides Echinodermata, compounds isolated from representatives of the phylum Porifera (marine sponges) also attract the interest of researchers working on the tuberculosis problem. Thus, it has been shown that alkaloids isolated from marine sponges can suppress Mtb growth[45].

To search for new non-azole inhibitors of CYP124 among natural compounds, we used standard P450 ligand screening methods (SPR analysis[46,47] and spectrophotometric titration [48,49]). Screening a library of 32 plant and marine compounds identified 9 new ligands for the active site of Mtb CYP124. *In silico* models of complexes were obtained for a number of them. Analysis of structurally similar (>0.4 Tanimoto) human endogenous metabolites from the KEGG database identified a group of 37 compounds, including immunoregulatory sterols and their precursors, some of which are already known as CYP124 ligands. The new CYP124 ligands we found may represent promising core structures for the development of compounds with higher affinity for CYP124. Further biochemical analysis in a reconstituted Mtb CYP124 system confirmed that some of the found compounds are new natural non-azole inhibitors of the enzyme. Among them, 25S)-5α-cholestane-3β,4β,6α,7α,8,15β,16β,26-octaol (15β-octaol) and henricioside Н_2_ (HD-4) acted as inhibitors forming long-lived complexes (τ1/2 of the CYP124/inhibitor complexes was 181 and 65 min, respectively). The obtained results expand knowledge about the structures of potential ligands and inhibitors of the pharmacologically significant Mtb cytochrome P450, representing a new potential target for tuberculosis pharmacotherapy.

## 2. MATERIALS AND METHODS

### 2.1 Reagents

HBS-N buffer (10 mM HEPES, 150 mM NaCl, pH 7.4), 10 mM sodium acetate buffers (pH 4.5, 5.0, and 5.5), a reagent kit for the covalent immobilisation of proteins via primary amino groups (1-ethyl-3-(3-dimethylaminopropyl)-carbodiimide-HCl (EDC), and N-hydroxysuccinimide (NHS)) were obtained from Cytiva (Marlborough, MA, USA).

5-cholesten-3β-ol-7-one was obtained from Steraloids Inc. (Newport, R.I., USA), NADPH – from Glentham Life Science (UK), isocitrate dehydrogenase – from Sigma Life Science (Oakville, ON, Canada), D-glucose-6-phosphate – from AppliChem GmbH (Darmstadt, Germany).

The remaining reagents were obtained from local suppliers.

### 2.2 Equipment

Protein–ligand interaction analysis was performed using SPR biosensors Biacore 8K and X100 (Cytiva, Marlborough, MA, USA). Refractive indexes of running buffers and experimental samples were matched with a refractometer RX-5000 (Atago, Saitama, Japan). Spectrophotometric titration was performed using a Cary 5000 UV–Vis NIR dual-beam spectrophotometer (Agilent Technologies, Santa Clara, CA). The HPLC-MS experiments were carried out with an Agilent HPLC 1200 instrument equipped with an Agilent Triple Quad 6410 mass spectrometer (Agilent Technologies, Santa Clara, CA, USA).

### 2.3 Cloning, Expression, and Purification of Proteins

Highly purified (>95% according to SDS-PAGE) preparations of recombinant proteins (CYP124, RubB, Arh1G18A) were carried out at the Institute of Bioorganic Chemistry of the National Academy of Sciences of Belarus using molecular cloning and heterologous expression in a bacterial system (*E. coli*) followed by metal affinity and ion exchange chromatography for protein purification. The plasmid for expression of *S. pombe* ferredoxin reductase Arh1 (mutant A18G) was kindly provided by Prof. Rita Bernhardt (Saarland University, Saarbrucken, Germany). The proteins (CYP124[22], RubB[50], Arh1G18A[51]) were cloned, expressed and purified as described previously.

### 2.4 Obtaining of Investigated Compounds

The studied compounds were isolated or obtained by chemical modifications at the G.B. Elyakov Pacific Institute of Bioorganic Chemistry, Far Eastern Branch of the Russian Academy of Sciences, similar to protocols published earlier [52–64]. The polar steroidal compounds (24*S*)-5α-cholestane-3β,4β,6β,8,15β,24-hexaol (HD-3) and henricioside Н_2_(HD-4) were isolated from the Far Eastern starfish *Henricia derjugini*[52]; spiculiferosides A–C were obtained from the Far Eastern starfish *Henricia leviuscula spiculifera*[53]; (24*E*)-5α-cholest-24-ene-3β,4β,6α,8,14,15α,26-heptaol 15-*O*-sulfate (А10) was isolated from the Vietnamese starfish *Archaster typicus*[54]; asterone (Agl1) was obtained by hydrolysis of the asterosaponin fraction of the Far Eastern starfish *Lethasterias nanimensis chelifera*[55]; (25*S*)-5α-cholestane-3β,4β,6α,7α,8,15β,16β,26-octaol (15β-octaol) was isolated from the Far Eastern starfish *Patiria pectinifera* as described in[56]; (25*S*)-5α-cholestane-3β,4β,6α,7α,8,15α,16β,26-octaol 3-О-sulfate (3-OSO_3_-octaol) and (25*S*)-5α-cholestane-3β,4β,6α,7α,8,15α,16β,26-octaol 3,26-О-disulfate (3,26-di-OSO_3_-octaol) were obtained by sulfation of (25*S*)-5α-cholestane-3β,4β,6α,7α,8,15α,16β,26-octaol (15α-octaol) isolated from the starfish *P. pectinifera* as described in[56]. The triterpene glycoside hemioedemoside A was isolated from the sea cucumber *Colochirus robustus*[57]. The isomalabaricane triterpenoids stellettins Q and R, andnor-isomalabaricanes jaspolide F, and globostelletin G were obtained from the Vietnamese marine sponge *Rhabdastrella globostellata* as described in[58,59]. The sponge was assigned to the genus *Stelletta* and then re-identified and reported as *R. globostellata* species[65]. The bibenzochromenone phanogracilin A (570-2) was isolated from the crinoid *Phanogenia gracilis* collected in the South China Sea[60]. The quinoid pigments echinochrome A, echinamine A, and echinamine B were isolated from the sea urchin *Scaphechinus mirabilis*[61], and spinochromes B, D and E were isolated from the sea urchin *Mesocentrotus nudus*[62]. The polyphenolic compounds retusin, maackiain, tectorigenin, liquiritigenin, medicarpin, formononetin, piceatannol, maackin, and (±)-3-hydroxyvestitone were isolated from *Maackia amurensis* as described in[63]. Luteolin 7,3′-disulfate was obtained from sea plants of the Zosteraceae family as described in[64].

Detailed protocols for compound isolation and purification are provided in the Supplementary Material (Section 1).

### 2.5 SPR Analysis

Real-time measurements of CYP124-ligand interactions were performed at 25 °C using CM5 optical chips coated with carboxymethyl dextran (CM-dextran). Sensorgrams were recorded as the change in the biosensor signal in resonance units (RUs) per unit of time (s). CYP124 was covalently immobilised onto the CM5 chip via amino groups. The carboxyl groups of CM-dextran were activated using a 1:1 v/v mixture of 0.4 M EDC and 0.1 M NHS for 7 min at a flow rate of 5 µL/min. CYP124 (20 μg/mL) in the 10 mM sodium acetate buffer (pH 5.0) was injected for 5 min at a flow rate of 5 μL/min. Next, the chip surface was washed with HBS-N buffer for 60 min. During interaction recording, a chip cell without immobilised CYP124 was used as the reference channel.

To screen the ability of natural compounds to bind to CYP124, analytes (50 μM) were injected over the surface of the chip with immobilised CYP124 for 10 minutes at a flow rate of 10 μL/min. Complex dissociation was recorded for 5 min at the same flow rate. Chip surface regeneration after each compound injection was performed by double injection of regenerating solution (2M NaCl, 0.5% CHAPS) for 20 s at a flow rate of 30 μL/min. Analysis was performed at 25°C. The resultant binding signal of the test compound with the target protein was recorded 5 minutes after the start of the dissociation phase of the formed protein-compound complex. The screening pass criterion was a registered signal amplitude **>**10 RU. Subsequently, visual analysis of the obtained sensorgrams was performed to prioritise the identified hit compounds.

To evaluate the half-life values of hit compound complexes with CYP124 (τ_1/2_), compound solutions (10-100 μM) were injected in “Multi-cycle” mode for further calculation of the equilibrium dissociation rate constant (k_off_). The injection time for each concentration was 210 s, the dissociation recording time after the end of the series of injections of the low molecular weight compound was 15 min, and the flow rate was 15 μL/min. The compounds’ stock solutions were prepared as 10 mM stock solutions in 100% dimethyl sulfoxide (DMSO). Experimental samples of compounds were prepared in HBS-N buffer at a concentration range of 10–100 μM and 1% DMSO. The same amount of solvent was added to the HBS-N running buffer to minimise bulk effects introduced by the difference between the refractive indexes of the running buffer and the experimental samples. If needed, the solvent concentration in the running buffer was corrected according to the equation:

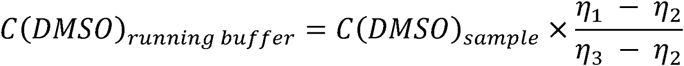

where *C(DMSO)_running buffer_* – DMSO final concentration in running buffer, *C(DMSO)_sample_*– DMSO concentration in experimental sample, *η*_1_– analysed sample refractive index, *η*_2_ – HBS-N buffer refractive index, *η*_3_ – HBS-N buffer containing the DMSO of the same concentration as experimental sample refractive index.

SPR sensorgrams were processed with Biacore X100 Evaluation Software Version 2.0.1 Plus Package (Cytiva, USA) and Biacore Insight Evaluation Software version 3.0.11.15423 (Cytiva, USA) using “1:1 dissociation”. The obtained k_off_ values were converted to τ_1/2_ using the formula:

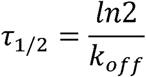

### 2.6 Ligand Binding Assay

Binding of ligands with the active site of CYP124 was carried out by spectrophotometric titration in 1 cm quartz cuvettes. Stock solutions of the ligands were prepared at a concentration of 10 mM in DMSO. The titration was repeated at least three times and ligand-binding constants (Kd_app_ values) were calculated using the equation as described previously as well as using the Hill equation [22].

### 2.7 Catalytic Activity Assays

Catalytic activity of CYP124 was reconstructed as described previously [23]. For evaluation of compounds’ inhibitory potentials, proteins (CYP124, Arh1G18A *S. pombe* and RubB *M. tuberculosis*) were preincubated with the substrate 7-keto-cholesterol and compounds were added at concentrations ranging from 10 to 100 µM.

### 2.8 Bioinformatics

#### 2.8.1 Prediction of Biological Activity Spectra

The web service PassOnline[66] was used to predict the spectra of biological activity for the investigated compounds (which showed the ability to inhibit CYP124 *in vitro*). SDF files with the structures of the hit compounds were used as input data for PassOnline. The outgoing information in text format was further analysed to identify the most significant possible effects to exclude potential toxicity.

#### 2.8.2 Search for Structures Similar to the Investigated Compounds Among Classical Human Endogenous Metabolites

The following algorithm was used to search among classical human endogenous metabolites for compounds that have similarity to the lead compounds from our dataset. First, a list of compounds from the KEGG database[67] exhibiting a Tanimoto similarity coefficient >0.4 to our target compounds (as of 30.05.25) was obtained. This cut-off is based on the fact that a Tanimoto coefficient value >0.4 is the minimum significant indicator of similarity for two low molecular weight compounds[68]. The obtained lists of compounds were analysed to exclude phytosterols and polyketides present in the human body as products of metabolism of dietary compounds. Modulators of the gut microbiota (e.g., bile acids) were also excluded.

#### 2.8.3 Modelling of Complexes

##### 2.8.3.1. Molecular Docking

Docking of compounds showing *in vitro* ability to inhibit the investigated cytochrome into the CYP124 active site was performed in the Flare 10.0.1 program (Cresset Group, UK). Files of 3D structures of CYP124 in various conformations were taken from the PDB site: 6T0G (complex with vitamin D3), 6T0F (complex with cholest-4-en-3-one), 6T0K (complex with carbethoxyhexyl imidazole). These crystal structures were described in detail in the works of Varaksa et al.[23]and Bukhdruker et al.[69]. For all structures, extraction of the ligand and other small molecules was performed, followed by energy minimisation. Structures of the investigated compounds were built based on data from the corresponding compound SDF files. Ligand docking was performed in “Normal” mode, calculation mode “Very accurate but slow”, without using a ligand template. Binding site search was conducted within the CYP124 active site cavity, including the substrate access channel and heme. Selection of ligand-protein binding poses was performed according to the following criteria: 1) the distance from the heme Fe to the nearest non-hydrogen atom of the ligand was no more than 4 Å (the value was taken considering that in the crystal structures used for docking, CYP124 ligands were at distances of 3.99 (6T0F), 4.00 (6T0G), and 2.1 (6T0K) Å); 2) the orientation of the ligand in the active site cavity did not cause steric conflicts.

##### 2.8.3.2. Molecular Dynamics

For molecular dynamic (MD) investigation of CYP124 interactions with HD-4 and 15β-octaol, molecular docking was performed using Molecular Operating Environment CCG version 2020.09 (MOE) (Montreal, Canada). The crystal structure of *M. tuberculosis* strain Rv2266 monooxygenase CYP124 complexed with cholest-4-en-3-one (PDB ID 6TOF) served as the receptor [23]. Ligands were prepared via MOE protonation modules. Flexible docking with “induced-fit” protocol ranked ligand poses, followed by force field refinement of the top 100 poses, GBVI/WSA dG rescoring, and detailed analysis of the best-ranked pose after solvation and energy minimisation.

Ligands under investigation were ranked by blind flexible docking with the “induced fit” protocol. Prior to docking, the force field of AMBER10:EHT and the implicit solvation model of the Reaction Field (R-field) were selected. Optimisation of both the protein’s protonation status and hydrogen bonding arrangements were carried out employing the Structure Preparation module at biologically relevant conditions—specifically, pH 7.4 and a temperature set at 300 Kelvin. During the flexible docking phase, initial ligand configurations were first prioritised based on their respective London dG energy values. This preliminary sorting was succeeded by detailed refinements involving the most promising 100 docked structures, which were subjected to additional fine-tuning using MD-based optimisation. These optimised structures were subsequently rescored with greater precision using the GBVI/WSA dG methodology. Ultimately, the single configuration achieving the highest rank in these assessments was preserved for subsequent analyses.To ensure the HD-4 and 15β-octaol best docking poses near CYP124 may be close to realistic ones (or stable) they were underwent to short-term MD simulations. The started system, solvated with about 19598 water molecules in 0.1M NaCl (18-19Na^+^ and 2-4 Cl^−^ depending on system). All-atomic MD simulations of 20 ns-long was carried out at constant pressure (1 atm) and temperature (300 K) with a time step of 2 fs. All-atomic force field Amber 10:EHT was used for proteins[70]; for water, TIP3P was used[71]. Solvent molecules were treated as rigid. All MD simulations including heating, equilibration as well as energy minimization steps were performed with MOE2019.01 CCG software package using cluster CCU “Far Eastern computing resource” FEB RAS (Vladivostok). The analysis of the noncovalent intermolecular interaction contributions to the free energy was performed by using the *IF-E* 6.0 SVL script tool [72], which enables calculation of the interaction energy on a per-residue basis (measured in kcal/mol). Negative energy values reflect favorable interactions between the ligand and target molecule, whereas positive values correspond to energetically unfavorable configurations. This computational tool has been successfully applied to interpret the ligand-receptor binding specificity in the active site of various adenosine receptor types [73–75].

## 3. RESULTS

The pathway of the work presented at Figure 2. A comprehensive approach to search for prototypes of potential CYP124 inhibitors was used and SPR screening, spectral titration, biochemical testing of the ability to inhibit CYP124 activity, as well as investigation of the “CYP124/inhibitor” complex half-life and IC50 and*in silico* modelling.

**Figure 1.**
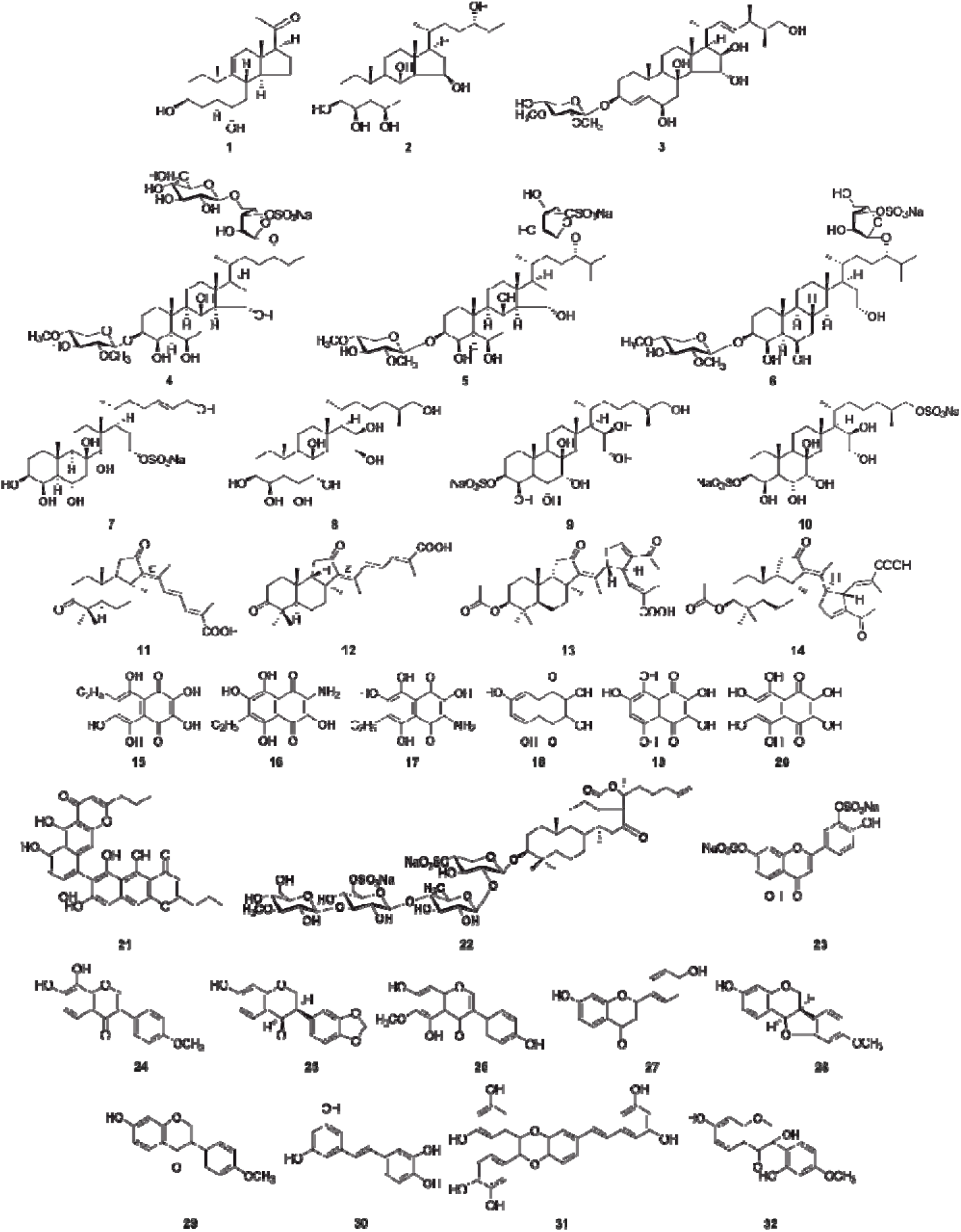
Structures of natural compounds. Asterone from the Far Eastern starfish *Lethasterias nanimensis chelifera* (Agl1) **(1)**; (24S)-5α-cholestane-3β,4β,6β,8,15β,24-hexaol (HD-3) **(2)** and Henricioside Н_2_(HD-4) **(3)** from the Far Eastern starfish *Henricia derjugini*; Spiculiferosides A **(4)**, B **(5)** and C **(6)** from the Far Eastern starfish *Henricia leviuscula spiculifera*; (24E)-5α-cholest-24-ene-3β,4β,6α,8,14,15α,26-heptaol 15-O-sulfate (А10) from the Vietnamese starfish *Archaster typicus* **(7)**; (25S)-5α-cholestane-3β,4β,6α,7α,8,15β,16β,26-octaol (15β-octaol) from the Far Eastern starfish *Patiria pectinifera* **(8)**; (25S)-5α-cholestane-3β,4β,6α,7α,8,15α,16β,26-octaol 3-О-sulfate (3-OSO -octaol) **(9)** and (25S)-5α-cholestane-3β,4β,6α,7α,8,15α,16β,26-octaol 3,26-О-disulfate (3,26-di-OSO -octaol) **(10)** obtained by sulfation of (25S)-5α-cholestane-3β,4β,6α,7α,8,15α,16β,26-octaol (15α-octaol) from the starfish *Patiria pectinifera*; Globostelletin G **(11)**, Jaspolide F **(1)** and Stellettins Q **(13)** and R **(14)** from marine sponge *Rhabdastrella globostellata*; Echinochrome A **(15)**, Echinamines A **(16)** and B **(17)** from the sea urchin *Scaphechinus mirabilis*; Spinochromes B **(18)**, D **(19)** and E **(20)** from the sea urchin *Mesocentrotus nudus*; Phanogracilin A (570-2) from the crinoid *Phanogenia gracilis* **(21)**; Hemioedemoside A from the sea cucumber *Colochirus robustus* **(22)**; Luteolin 7,3′-disulfate from the sea plant of Zosteraceae genus **(23)**; Retusin **(24)**, Maackiain **(25)**, Tectorigenin **(26)**, Liquiritigenin **(27)**, Medicarpin **(28)**, Formononetin **(29)**, Piceatannol **(30)**, Maackin **(31)** and (±)-3-hydroxyvestitone **(32)** from the Far Eastern endemic plant *Maackia amurensis*.

**Figure 2.**
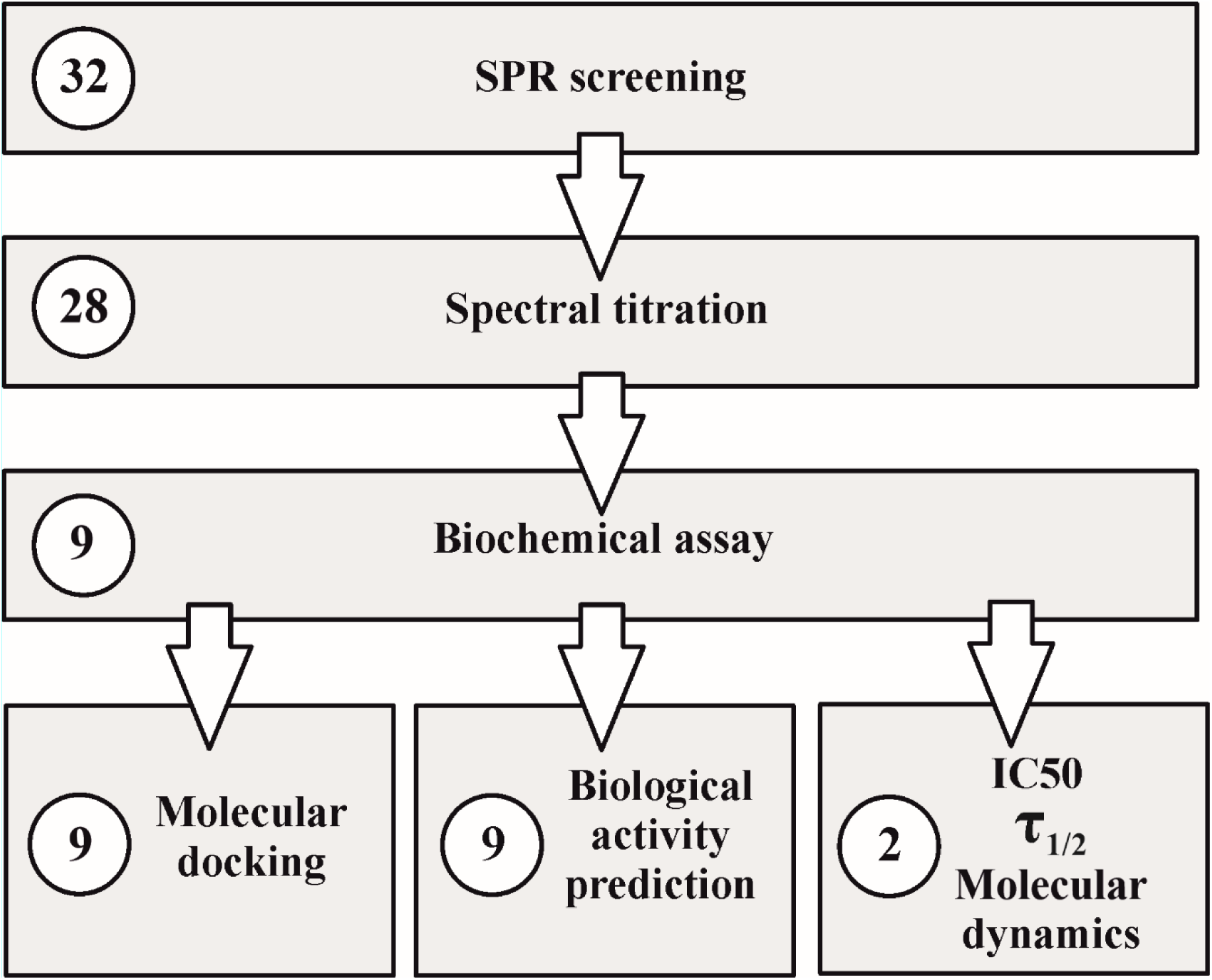
The pathway of the work presented. The number of compounds used in the given stage of analysis presented in circles.

### 3.1 Screening of the Ligand Sample for Binding Ability to CYP124

To select ligands from the investigated sample consisting of 32 natural compounds, screening was performed on an SPR biosensor. This method assesses the compound’s ability to bind to the CYP124 molecule. To evaluate the interaction of compounds with the CYP124 active site, spectral titration was performed. Results are presented in **Table 1**. 4 compounds failed the SPR screening. For all compounds showing a positive result in spectral titration, a Type I (maximum absorption at wavelengths of 385–390 nm and minimum absorption at 420 nm)[76]spectral response was characteristic (spectra are provided in Supplementary, **Table S1**). Among the compounds that passed SPR screening, a spectral response allowing reliable calculation of Kd was registered for 9.These 9 compounds were subsequently selected for testing inhibition of CYP124 enzymatic activity *in vitro*. For the remaining compounds, which were characterized by a weak spectral response or its absence, accurate determination of the Kd value was not feasible.

**Table 1.** Interaction of natural compounds with CYP124.

| № | Compound | SPR-screening | Spectral titration<br>/ Kd spectral for<br>++ (uM) | CYP124<br>inhibition, % |
| --- | --- | --- | --- | --- |
| **starfish *Lethasterias nanimensis chelifera*,**<br>**hydrolysis** |  |  |  |  |
| 1 | Asterone (Agl) | + | + | n.d. |
| **starfish *Henricia derjugini*** |  |  |  |  |
| 2 | (24*S*)-5α-Cholestane-3β,4β,6β,8,15β,24-hexaol (HD-3) | + | + | n.d. |
| 3 | Henricioside H<sub>2</sub> (HD-4) | + | ++ / 4.06 (type I) | 31 |
| **starfish *Henricia leviuscula spiculifera*** |  |  |  |  |
| 4 | Spiculiferoside A | + | + | n.d. |
| 5 | Spiculiferoside B | + | ++ / 4.08 (type I) | 5 |
| 6 | Spiculiferoside C | + | ++ / 7.61 (type I) | <1 |
| **starfish *Archaster typicus*** |  |  |  |  |
| 7 | (24*E*)-5α-Cholest-24-ene-3β,4β,6α,8,14,15α,26-heptaol 15-*O*-sulfate (A10) | + | + | n.d. |
| **starfish *Patiria pectinifera*** |  |  |  |  |
| 8 | (25*S*)-5α-Cholestane-3β,4β,6α,7α,8,15β,16β,26-octaol (15β - octaol) | + | ++ / 10.37 (type I) | 38 |
| **starfish *Patiria pectinifera*, sulfation** |  |  |  |  |
| 9 | (25*S*)-5α-Cholestane-3β,4β,6α,7α,8,15α,16β,26-octaol-3-*O*-sulfate (3-OSO<sub>3</sub>-octaol) | + | ++ / 15.62 (type I) | 23 |
| 10 | (25*S*)-5α-Cholestane-3β,4β,6α,7α,8,15α,16β,26-octaol-3,26-*O*-disulfate (3,26-di-OSO<sub>3</sub>-octaol) | + | ++ / 20.22 (type I) | 22 |
| **marine sponge *Rhabdastrella globostellata*** |  |  |  |  |
| 11 | Globostelletin G | + | - | n.d. |
| 12 | Jaspolide F | + | - | n.d. |
| 13 | Stellettin Q | + | ++ / 8.25 (type I) | 2 |
| 14 | Stellettin R | + | - | n.d. |
| **sea urchin *Scaphechinus mirabilis*** |  |  |  |  |
| 15 | Echinochrome A | + | - | n.d. |
| 16 | Echinamine A | + | - | n.d. |
| 17 | Echinamine B | + | - | n.d. |
| **sea urchin *Mesocentrotus nudus*** |  |  |  |  |
| 18 | Spinochrome B | + | - | n.d. |
| 19 | Spinochrome D | + | - | n.d. |
| 20 | Spinochrome E | + | - | n.d. |
| **crinoid *Phanogenia gracilis*** |  |  |  |  |
| 21 | Phanogracilin A (570-2) | + | - | n.d. |
| **sea cucumber *Colochirus robustus*** |  |  |  |  |
| 22 | Hemioedemoside A | - | - | n.d. |
| **sea plant of *Zosteraceae* family** |  |  |  |  |
| 23 | Luteolin 7,3'-disulfate | + | - | n.d. |
| **Far Eastern endemic plant *Maackia amurensis*** |  |  |  |  |
| 24 | Retusin | + | - | n.d. |
| 25 | Maackiain | + | ++ / 140.90 (type I) | 14 |
| 26 | Tektorigenin | - | - | n.d. |
| 27 | Liquiritigenin | + | - | n.d. |
| 28 | Medicarpin | + | ++ / 22.55 (type I) | 17 |
| 29 | Formononetin | - | - | n.d. |
| 30 | Piceatannol | + | - | n.d. |
| 31 | Maackin | + | - | n.d. |
| 32 | (±)-3-hydroxyvestitone | - | - | n.d. |

SPR Screening: Binding was considered positive (+) if the response exceeded 15 RU after compound injection; a lack of binding is denoted as (-). Spectral Titration: Interactions were classified as: (++) strong binding with a characteristic spectral response and a reliably determined Kd value; (+) weak binding with an unreliable Kd estimation; (±) interaction with the heme iron (confirmed by a Soret band shift to 417–420 nm) in the absence of a characteristic spectral signature; (-) no spectral response. CYP124 Inhibition Test: The percentage of enzymatic inhibition was measured at a 50 µM compound concentration. "n.d." indicates that the inhibition was not determined for that compound.

### 3.2 In Vitro Test for Inhibition of CYP124 Enzymatic Activity

An *in vitro* assay of the CYP124-catalysed oxidation reaction of 7-keto-cholesterol in the presence of the 9 priority compounds selected at the screening stage, at a concentration of 50 μM of the test compound, was performed (**Table** 1). The reference inhibitor compound N′-hydroxy-N-(4-isopentyl-2-methylphenyl)formimidamide (cpd5’) was used. The action of cpd5’ as a CYP124 inhibitor was previously demonstrated in the work of Tatsiana Varaksa et al. [23]. At a cpd5’ concentration of 50 μM, 80% inhibition of CYP124 activity was achieved.

For compounds showing CYP124 inhibition >30%, IC50 was determined by titration in the concentration range 10–100 μM). It was found that for 15β-octaol IC50 ≈86 μM. For HD-4, IC50 >100 μM can only be stated. Cpd5’ showed an IC50 value <10 μM in this experiment.

### 3.3 Evaluation of CYP124-Ligand Complex Half-Life Times

Using the SPR biosensor, k_off_ values for complexes of HD-4 and 15β-octaol with CYP124 were evaluated, allowing calculation of τ_1/2_. It was established that the τ_1/2_values for complexes of CYP124 with 15β-octaol and HD-4 were 181 and 65 minutes, respectively. The sensorgrams of CYP124-compounds interactions are shown in Supplementary (**Figures S1-2**).

### 3.4. Prediction of Biological Activity Spectra for Hit Compounds

The natural molecules we studied can be considered as scaffolds for creating higher affinity and stronger inhibitors of CYP124. Accordingly, the question of whether the compounds we investigated have effects on the human body that may be associated with potential toxicity is relevant. As a result, the spectrum of potential pharmacological effects was predicted for the 9 compounds that gave the most characteristic spectral response (**Table** 1), presented in **Table S2** (only effects for which Pa (probability “to be active”) >0.7 were considered). Among the predicted effects, the 10 most potentially dangerous can be distinguished: immunosuppressive activity, antineoplastic activity (possible cytotoxicity), JAK2 expression inhibitor (possible pancytopenia), glyceryl-ether monooxygenase inhibitor (possible demyelination, retinopathies), 17β-HSD3 inhibitors (possible virilisation, osteoporosis), prostaglandin-E2 reductase inhibitors (possible gastrointestinal ulcers), sterol Δ14-reductase inhibitors (adrenal insufficiency, teratogenicity), testosterone 17β-dehydrogenase inhibitor (testosterone synthesis disorders), bilirubin oxidase inhibitors (jaundice, hepatitis), caspase 3 stimulant (neurotoxicity, hepatotoxicity, cardiotoxicity).

### 3.5 Search for Classical Endogenous Metabolites Similar to the Investigated Natural Compounds

The obtained list of 37 human endogenous classical metabolites exhibiting similarity to the lead compounds in our dataset was additionally filtered by the criteria “involvement in immunity or MTb-induced infectious process” or “presence of aliphatic chain at C17”. The first criterion was chosen based on considerations of Mtb’s influence on the course of human defence reactions through the metabolism of some steroids[23]. The second criterion, the presence of an aliphatic chain at the C17 atom of the steroid nucleus, is due to the fact that CYP124 tightly binds and hydroxylates these substrates at the chemically disfavoured ω-position[20].

As a result, a group of 37 compounds was formed (**Table** 2). These 37 compounds were divided into three groups: 1) those having a direct relationship to the regulation of immune processes; 2) those not directly related to immune reactions but possessing an aliphatic carbon chain at the C17 position; 3) compounds with questionable relationship with CYP124 due to the absence of the aliphatic chain at C17.

**Table 2.**
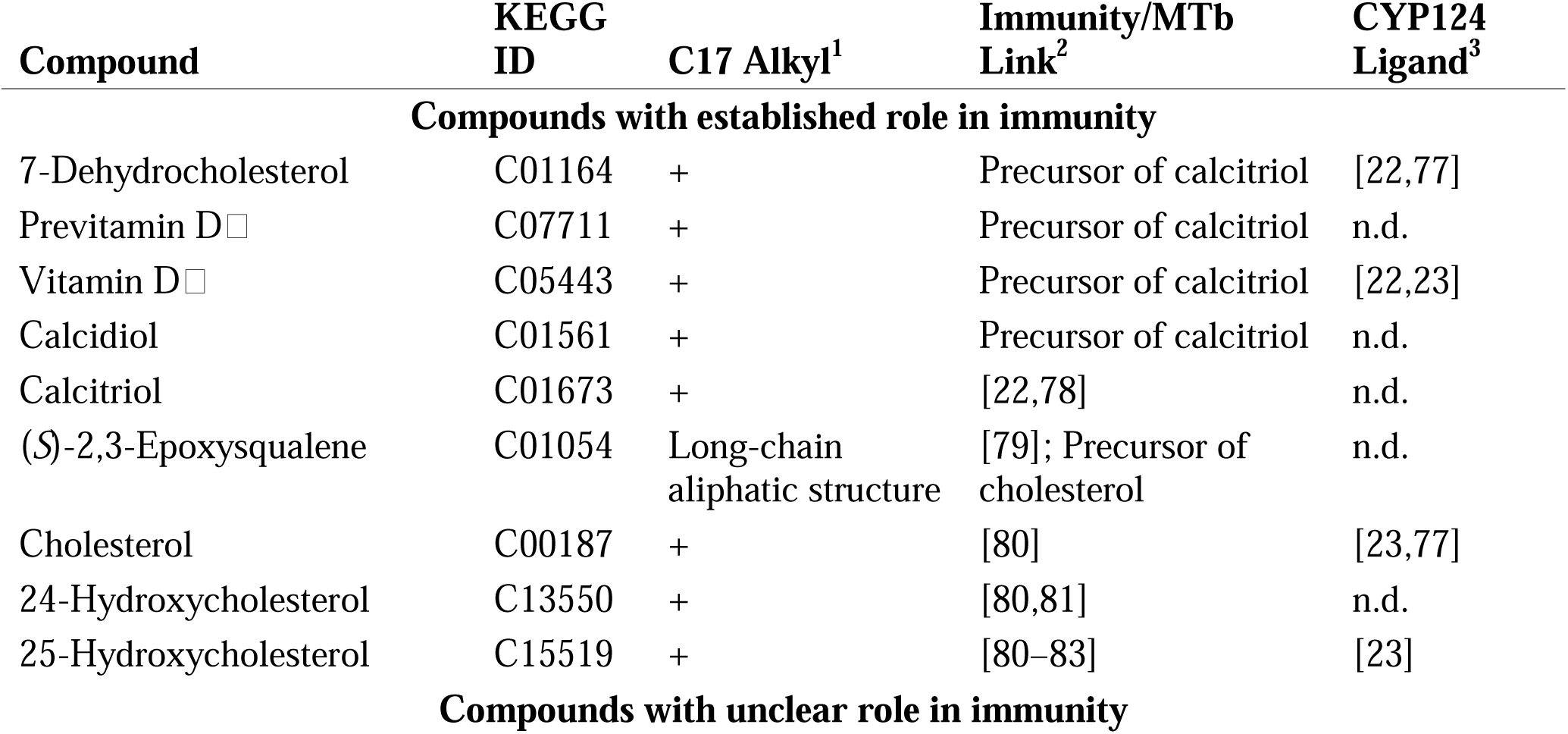
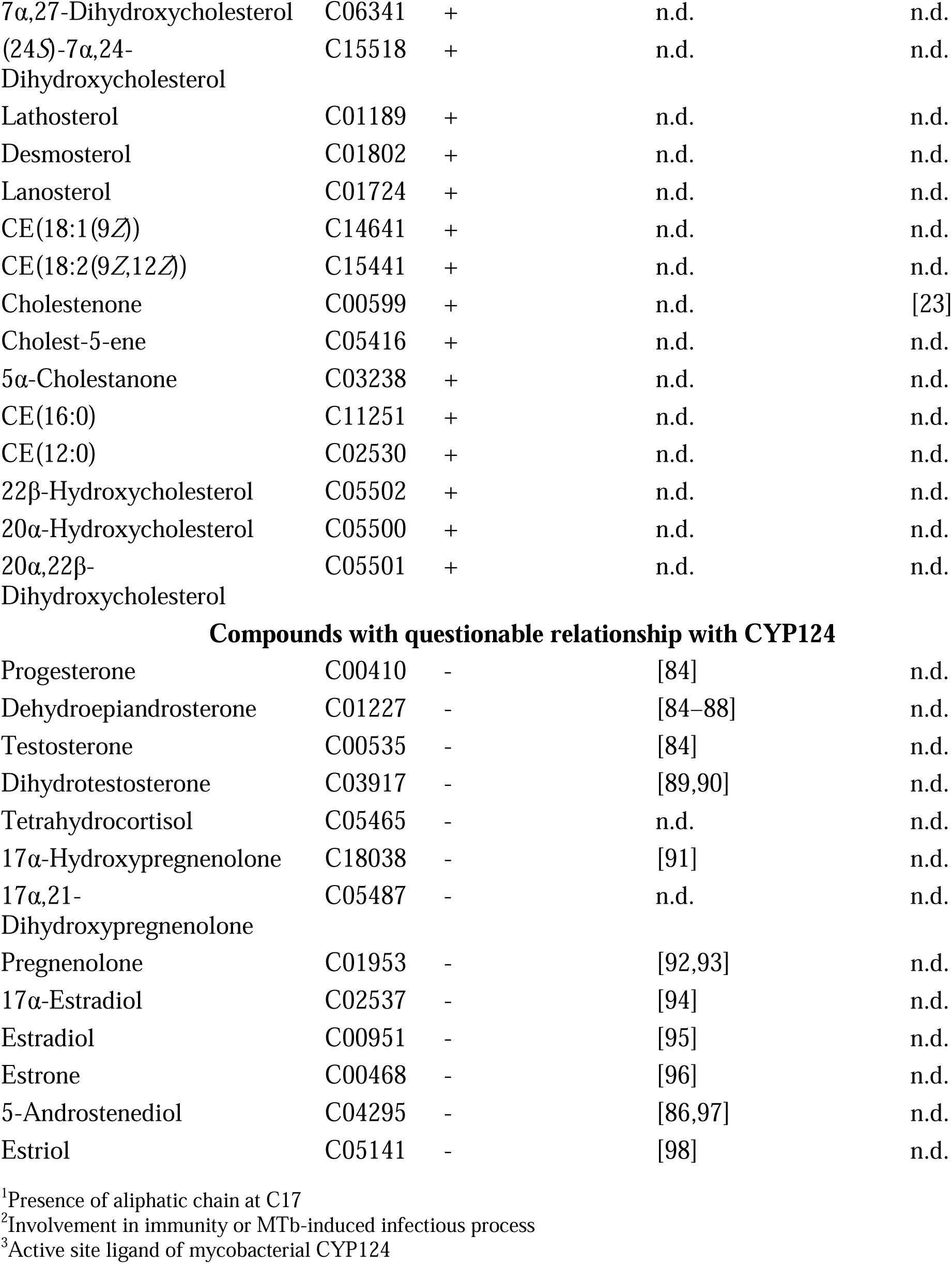
Lists of endogenous primary metabolites in humans exhibiting structural similarity (Tanimoto score >0.4) to the lead compounds in our dataset.

Presence of aliphatic chain at C17: “+” – there is the aliphatic chain at C17, “-” – there is no aliphatic chain at C17. n.d. – no data were found.

### 3.6 Modelling of CYP124-Ligand Complexes

Although the investigated ligands exhibit considerable structural diversity, we utilized *in silico* analysis to delineate the common binding features dictated by the CYP124 active site architecture. The primary objective was to identify the conserved hydrophobic core and key residues crucial for accommodating a broad spectrum of potential ligands. The identification of this shared pharmacophore serves to discriminate fundamental anchoring interactions from those that are ligand-specific. This approach thereby strengthens the interpretation of our experimental data and establishes a foundation for the rational design of future inhibitors.

#### 3.6.1 Molecular Docking

Molecular docking of the 9 ligands showing the best results in spectral titration into the active site of the CYP124 molecule was performed in the Flare 10.0.1 program. CYP124 crystals with different substrates (cholestenone, vitamin D3) and an inhibitor (carbethoxyhexyl imidazole) were used for docking. The use of different crystals was due to the significant conformational mobility of the CYP124 active site cavity, the configuration of which is significantly influenced by the structure of the bound compound[22,23,77]. As a result, using our docking parameters, it was not possible to obtain models that agreed with the spectral titration results and satisfied the selected selection criteria, except for 8 models (out of 27 possible protein-ligand combinations), which were deemed satisfactory (**Table 3**). When analysing the docking results, the main focus was on finding common CYP124 amino acids interacting with ligands in the models and crystal structures. This analysis was performed in the Flare program using the “Interaction map” tool. The obtained models of complexes of the investigated ligands with Mtb CYP124 are presented in Supplementary (**Figures S3-10**).

**Table 3.**
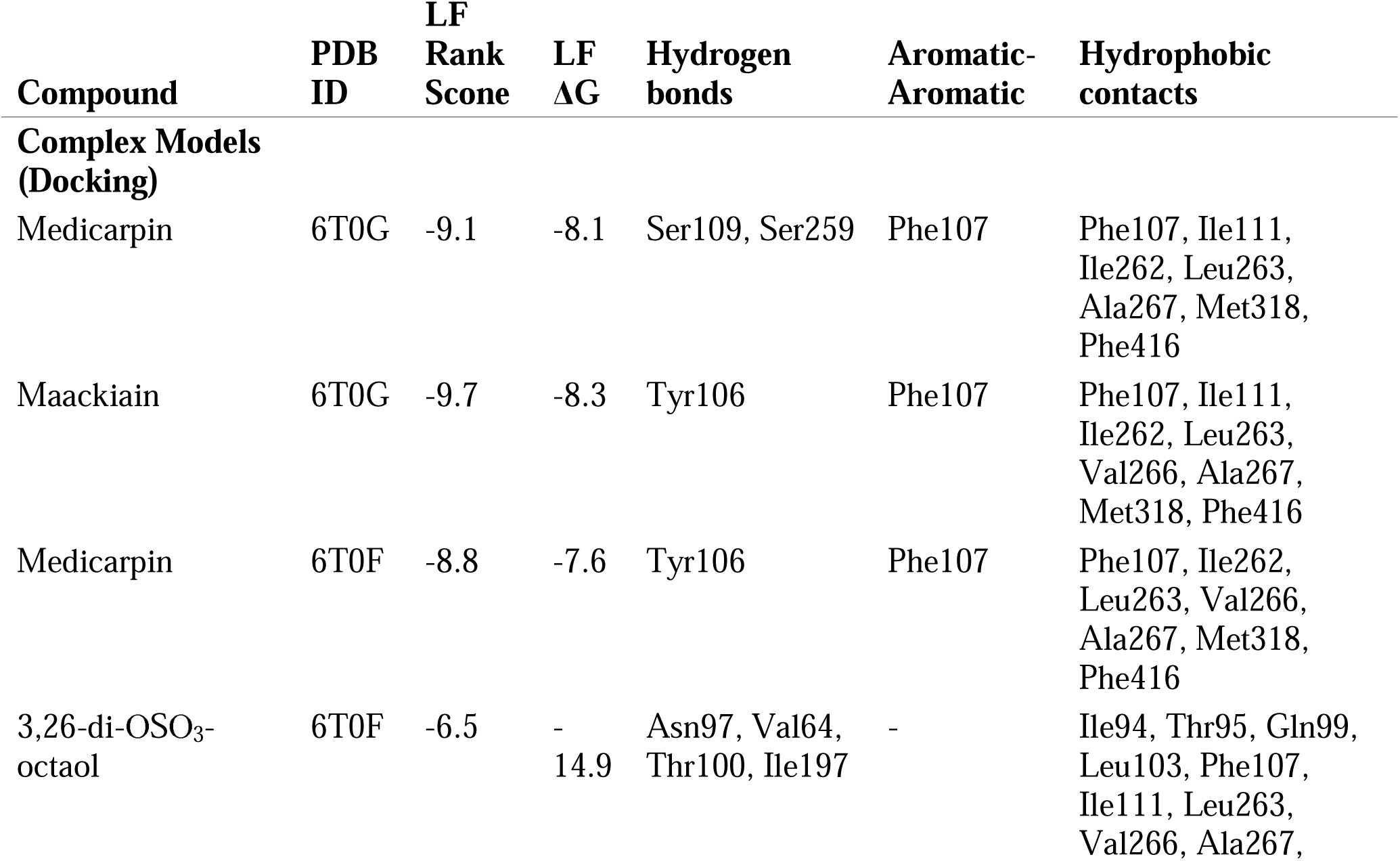
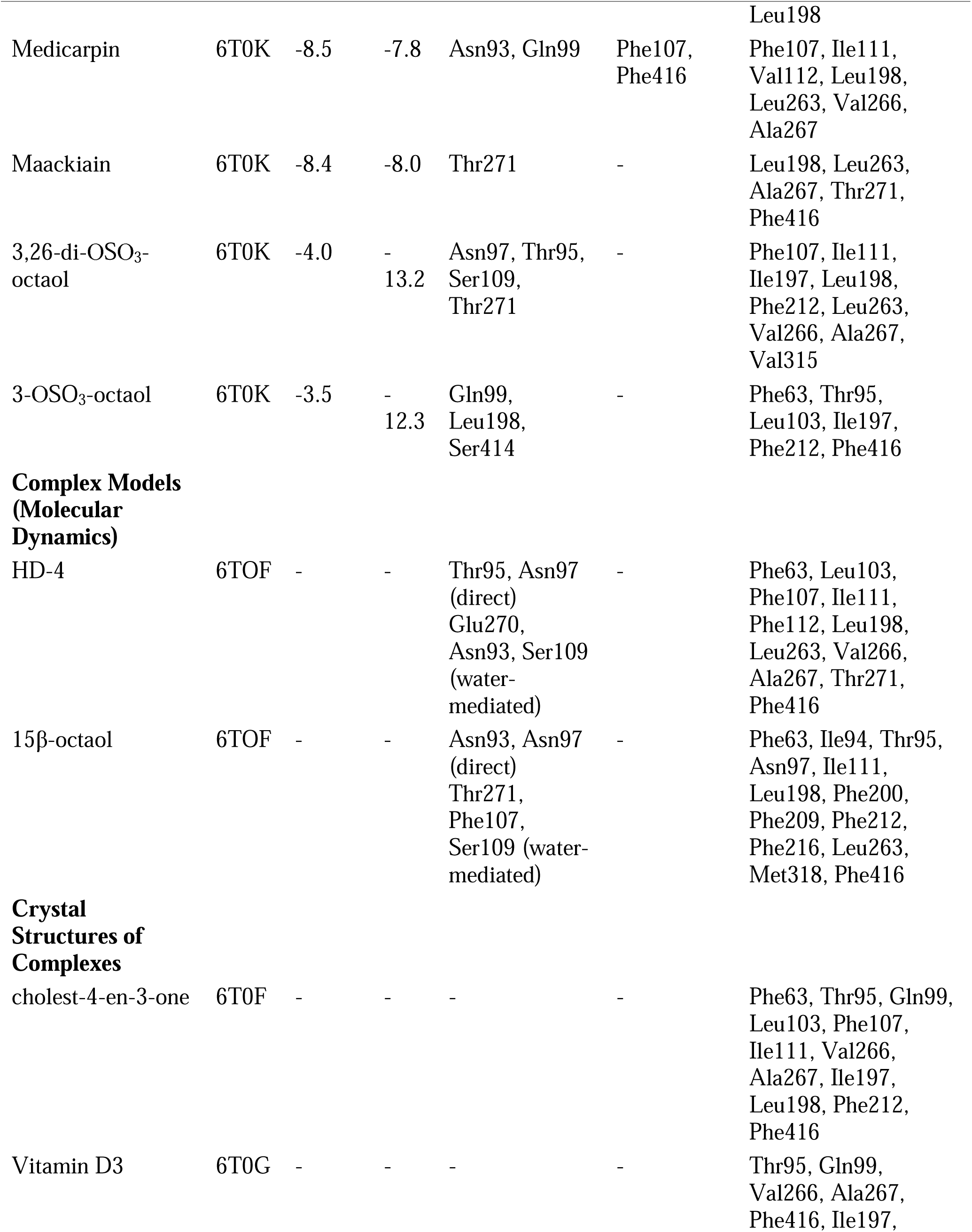
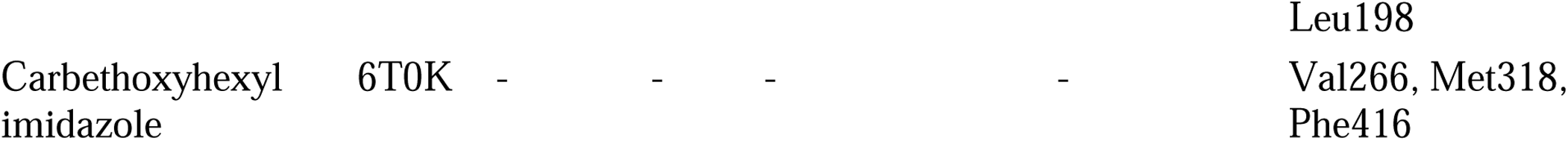
Parameters of models and crystal structures of compound complexes with CYP124.

#### 3.6.2 Molecular docking improved with short-term Molecular Dynamics simulations

Both HD-4 and 15β-octaol occupied the CYP124 binding pocket with significant overlapping with the cholest-4-en-3-one (cholestenone) binding site which serves as the cytochrome substrate [23] **(Figure 3)**. Optimized structures of complexes as well as the topology data are available via ZENODO repository (DOI: 10.5281/zenodo.17693346).

**Figure 3.**
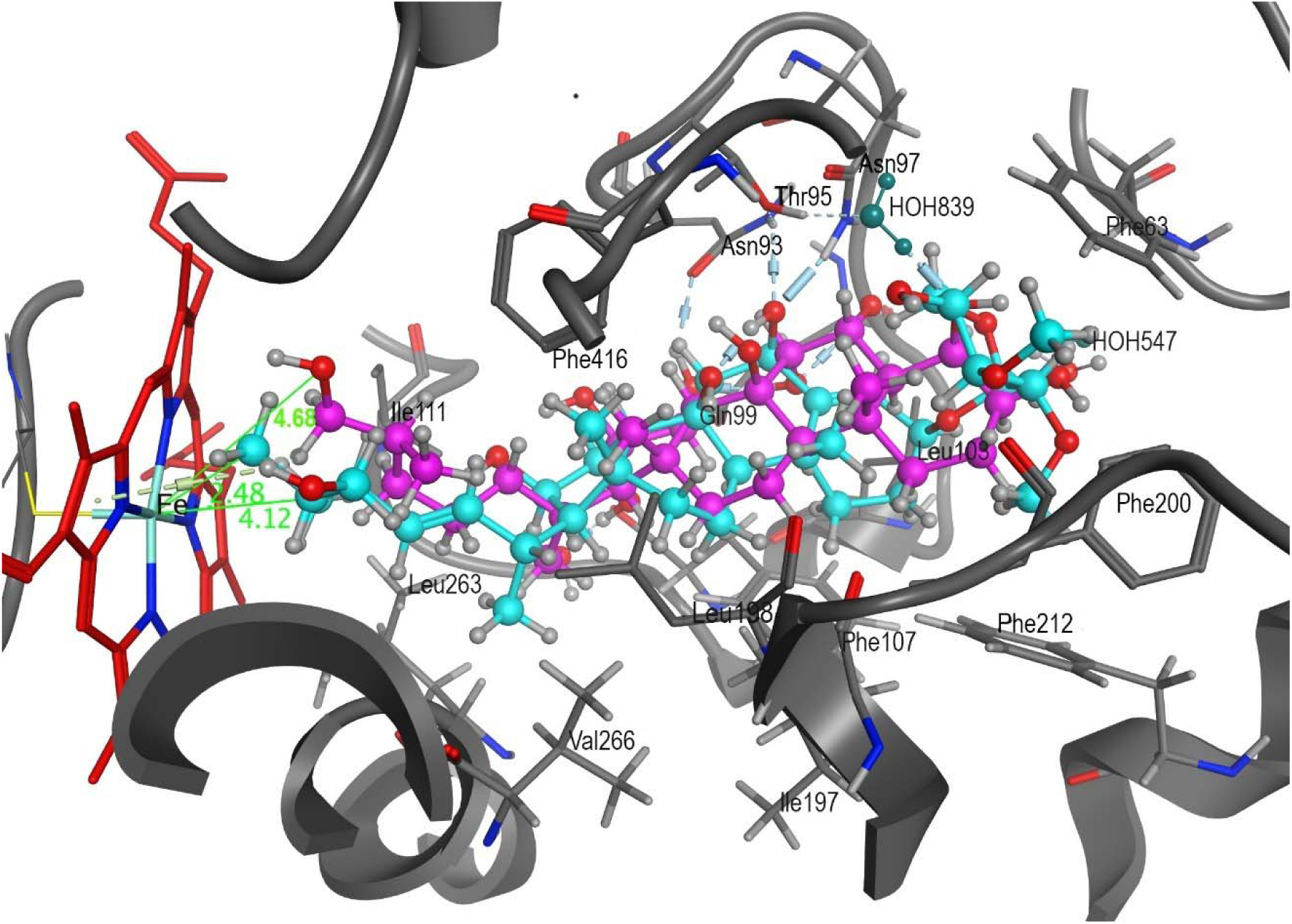
Diagram of the probable complex structures formed by Cyp124 with HD-4 and 15β-octaol. The receptor is depicted as a cartoon with the cholest-4-en-3-one (cholestenone) binding site residues represented as sticks (grey), with heme shown in stick representation (red), HD-4 (cyan) and 15β-octaol (magenta) are displayed as ball-and-stick model.

In the HD-4 complex, methyl groups 24-Me and 25-Me located at 4.12 Å and 2.48 Å, respectively, from the heme iron were engaged in dual H-π interactions (-1.3 kcal/mol total) and hydrophobic contacts (-6.9 kcal/mol). The carbohydrate chain formed H-π interactions with the aromatic rings of Phe63 (-1.0 kcal/mol), Phe112 (-0.3 kcal/mol), and the steroidal A-ring with Phe107 (-0.5 kcal/mol) (**Figure S11**). The 2,3-di-OMe-xylopyranose moiety exhibited hydrophobic stabilisation with Phe63 (-5.39 kcal/mol), Phe112 (-2.69 kcal/mol), Phe107 (-5.82 kcal/mol), and Leu103 (-2.15 kcal/mol). The HD-4 side chain formed hydrophobic clusters with side chains of Ile111, Val266, Thr271, Ala267, and Leu263 (-2.95 to -2.12 kcal/mol each) deep in the active site. The ligand core engaged hydrophobically with Phe416 (-2.19 kcal/mol), Leu198 (-1.39 kcal/mol), Thr95 (-2.47 kcal/mol), and Asn97 (-0.92 kcal/mol), while its 6β-hydroxy group acted as an H-bond acceptor with Thr95 and Asn97. Non-covalent intermolecular interactions analysis within this complex allowed us to identify the role of water molecules in HD-4 orientation in the Cyp124 binding site. Water-mediated interactions bridged the 26-hydroxy group to Glu270 (**Figure S11**) and connected the 15α-OH/3O-Me/C4-OH groups of the xylopyranose moiety to Ser109/Asn93/Asn97.

The 15β-octaol 26-OH group was positioned 4.68 Å from the heme iron (**Figure 3**). Hydrophobic interactions dominated complex formation: the ligand core with Phe212, Phe200, Phe63, Thr95, Asn97, Phe209, and Phe107 (from -4.13 to -2.10 kcal/mol), and the side chain with Leu198, Ile111, Leu263, Phe216, Ile94, and Met318. H-bonds between 8-OH/Asn93 (-2.6 kcal/mol) and 4β-OH/Asn97 (-2.3 kcal/mol) stabilised positioning (**Figure S12**). The Phe212 aromatic ring formed H-π contacts with the steroidal A-ring of the ligand. Water molecules facilitated interactions between Phe107 and Ser109 and the 16β-OH group via H-bonds. Hydrophobic/H-π interactions involving C25/26-OH and heme contributed -1.57 kcal/mol, with the 26-OH forming a water-mediated H-bond to Thr271.

An assessment of the interaction energy of Cyp124 with the solvent as well as with water molecules and ligand fragments involved in complexation occurring before and after binding revealed a gain in energy upon the complex formation with HD-4 and 15β-octaol of -32.85 kcal/mol and -10.10 kcal/mol, respectively. The computational results evidenced the compound HD-4 to be a more closely bound to CYP124 and is characterized by a lower Kd spectral value of 4.06 uM, derived from spectral titration experiments (**Table 1**), compared with that of 15β-octaol (Kd spectral of 10.37 uM), which has a less favorable binding energy value.

Divergent heme interactions were observed: HD-4 formed an H-bond with Glu373 (-7.7 kcal/mol; **Figure S13**), while 15β-octaol engaged in weaker H-bonds with Met318 (-0.30 kcal/mol) and Arg122 (-0.70 kcal/mol), plus ionic interactions with Arg122 (**Figure S14**). Conserved heme contacts (Ala267, Phe301, Glu370) occurred in both complexes.

Summary data on CYP124 amino acids involved in forming hydrogen bonds and hydrophobic contacts with ligands, according to MD models, are presented in **Table 3**.

## 4. DISCUSSION

Despite progress in treating bacterial infections, tuberculosis remains a global threat, with therapy for MDR-TB presenting particular complexity. Consequently, the search for chemical compounds that can act on new promising Mtb target proteins is relevant. Among such targets, cytochrome P450s have recently been highlighted[4,5]. CYP124 Mtb is considered one promising for pharmacological intervention; it is not vitally essential but plays a critically important role in virulence manifestation, particularly by being able to metabolise host immunoactive sterols [23]. CYP124 has wide substrate specificity [23,99]. As with many other cytochromes P450, minor changes in the structure of the molecule result in altered substrate binding to CYP124. These facts make the search for CYP124 Mtb inhibitors highly relevant. Our working hypothesis was that some representatives of the phylum Echinodermata*, genera Maackia* and *Zostera*might contain chemical compounds capable of binding to the active site of Mtb CYP124 and serving as structural bases for new inhibitors of this enzyme.

We applied a comprehensive approach (**Figure 2**), to search for prototypes of potential CYP124 inhibitors within a library of compounds isolated at the Pacific Institute of Bioorganic Chemistry. The pipeline included SPR screening, spectral titration, biochemical testing of the ability to inhibit CYP124 activity, as well as investigation of the “CYP124/inhibitor” complex half-life and IC50. Using *in silico* methods, models of lead compound complexes with CYP124 were obtained, their possible spectrum of biological activity was predicted, and a search for similar compounds among human endogenous metabolites was performed.

The first stage of the work was screening our library of 32 compounds isolated from natural sources on an SPR biosensor. As a result, 28 compounds showed the ability to interact with CYP124 (**Table** 1). Although SPR screening confirms the fact of interaction between the investigated ligand and the cytochrome, it does not provide information on the binding site location. SPR cannot establish whether ligands bind in the active site and interact with the heme iron, a factor important for a potential inhibitor. Therefore, at the next stage, the possibility of binding to the heme iron was evaluated for each of the 28 compounds using spectral titration. As a result, 9 compounds were selected for which a reliable Type I spectral response was obtained, allowing calculation of Kd (**Table 1**). A Type I spectral response indicates that the ligand can interact with the heme iron of the CYP, displacing a water molecule [100]. Thus, at this stage, we obtained a list of compounds capable of binding within the CYP124 active site cavity. Next, a biochemical test was performed to determine the ability of the selected 9 compounds to inhibit the enzymatic activity of CYP124 (model reaction of 7-keto-cholesterol oxidation) *in vitro* (**Table 1**). Despite relatively similar dissociation equilibrium constant (Kd) values, the ability of the compounds to inhibit CYP124 activity varied significantly.This observed effect may be attributed not only to the compounds’ interaction with the enzyme’s active site but also to other contributing factors. These may include binding to allosteric regulatory sites, direct interaction with the redox partners themselves, or interference at the protein-protein interface between the cytochrome P450 and its redox partner. For the two compounds showing the most significant inhibition at their constant concentration of 50 μM, 15β-octaol and HD-4, a series of additional studies were performed to obtain more information about their potential as scaffold structures for designing CYP124 inhibitors. Thus, to assess the potency of these inhibitors, we obtained IC50 values for the CYP124 model reaction. It was shown that 15β-octaol and HD-4 exhibited weaker inhibitory activity compared to the reference inhibitor Cpd5’. Another characteristic for evaluating the potential of chemical compounds as enzyme function inhibitors is the half-life of the protein-ligand complexes (τ_1/2_). It is currently considered that it is the lifetime (or residence time) of the binary drug–target complex that determines the extent and duration of drug pharmacodynamics[101]. We obtained τ_1/2_ values for 15β-octaol and HD-4 by SPR, amounting to 181 and 65 minutes, respectively. This is consistent with the described lifetime of FDA-approved drugs[102]. Also, it’s comparable to the half-life values of CYP3A4 complexes with azole inhibitors, ketoconazole and itraconazole, obtained by SPR [46]. Although 15β-octaol and HD-4 did not show low IC50 values, their high τ_1/2_ (181 and 65 min, respectively) indicate potential as core structures for creating new CYP124 inhibitors through their optimisation. This suggestion is grounded in the fact that the pharmacological relevance of a long residence time extends beyond the instantaneous affinity measured in a closed *in vitro* system. Under dynamic of the *in vivo* conditions, where drug concentrations fluctuate, the half-life of the drug-target complex is a critical determinant of efficacy and tends to correlate with it [103,104]. A prolonged τ1/2 ensures sustained target coverage even after systemic concentrations of the unbound drug decline below the IC50 or Kd value [103]. Moreover, efforts to improve the thermodynamic affinity and therefore lowering the IC50 of a drug may actually reduce in vivo efficacy [103].

However, we decided not to narrow our focus only to 15β-octaol and HD-4. It is advisable to consider all 9 compounds, as they are capable of being ligands of the CYP124 active site and their structures can be used as prototypes for potential inhibitors. The toxicity of the base structures is an important criterion for assessing the potential of structures for their further use as the basis for potential inhibitors. However, considering potential lead compounds must account not only for their potential to modulate the target protein’s function but also for probable toxic effects, because high toxicity can completely negate all efforts in developing a future drug[105]. Therefore, for a more balanced assessment of our 9 compounds, we predicted their spectra of potential biological activities using the PassOnline web service. As can be seen from **Table S2**, a wide list of potential biological activities was predicted for our sample of 9 ligands. We analysed the predicted biological activities and selected 10 from them that could be most dangerous in terms of toxicity manifestation: immunosuppressive activity, antineoplastic activity, JAK2 expression inhibitor, glyceryl-ether monooxygenase inhibitor, 17β-HSD3 inhibitors, prostaglandin-E2 reductase inhibitors, sterol Δ14-reductase inhibitors, testosterone 17β-dehydrogenase inhibitor, bilirubin oxidase inhibitors, caspase 3 stimulant. Among these effects, antineoplastic and immunosuppressive activities occur most frequently. The prediction of immunosuppressive activity is unfavourable, as this effect may be associated with toxic effects on immune system cells[106,107], and also cause disturbances in immune system function[106]. The predicted antineoplastic activity suggests that the compound could potentially be toxic to actively dividing cells (e.g., mucous membranes, haematopoietic tissues, etc.) [108]. For the compounds showing the greatest potential as prototypes for CYP124 inhibitors, 15β-octaol and HD-4, PassOnline predicted quite a few biological activities, including the most dangerous in terms of toxicity manifestation (**Table S2**). Thus, when using these structures as scaffold structures for potential CYP124 inhibitors, the possibility of such toxic effects must be considered. Special attention will also need to be paid to *in vivo* toxicity tests, as the toxic effects we mentioned are only predictions. However, in terms of the least number of predicted biological activities, stellettin Q and 3-OSO_3_-octaol can be considered priorities, for which two and three biological activities were predicted, respectively (**Table S2**). And if stellettin Q demonstrated extremely low inhibitory potential (2% inhibition of CYP124 activity in the model system), 3-OSO_3_-octaol can be considered more promising (**Table** 1). However, a possible antineoplastic effect was predicted for stellettin Q, which should be considered when working on optimising this compound’s structure. Among the biological activities predicted for 3-OSO_3_-octaol, the ability to inhibit two enzymes can be highlighted: Alkylacetylglycerophosphatase and Benzoate-CoA ligase. Alkylacetylglycerophosphatase inhibitor activity may be associated, for example, with impaired synthesis of platelet-activating factor (PAF)[109]. Interaction of PAF with its receptors results in the inhibition of cyclic AMP formation, mobilisation of intracellular Ca^2+^, and the activation of mitogen-activated protein kinases, leading to multiple biological effects [110], including disruption of platelet activation processes and immune reactions. The target for the other predicted activity of 3-OSO_3_-octaol, benzoate-CoA ligase, is characteristic of plants and bacteria[111,112]. Thus, in terms of potential for further structure modifications, 15β-octaol and HD-4 can be considered as the most potent, and also 3-OSO_3_-octaol, as having a small number of predicted biological activities with relatively good potency as a CYP124 inhibitor. It should be emphasised that all the biological activities we discuss are predictions, so they should be validated on *in vivo* models, and it is important to consider them in structure optimisation work.

In addition to considering compounds from our investigated sample as structure-scaffolds for new potential CYP124 inhibitors, we assessed which metabolites in the human body might also be ligands of this cytochrome’s active site. As a result of analysing lists of classical human endogenous metabolites obtained from the KEGG database, a group of 37 compounds was identified from the entire set (similarity to our 9 ligands >0.4 Tanimoto) (**Table 2**).

Analysis of the found literature allows us to propose the hypothesis that CYP124 *M. tuberculosis* may serve as a key link connecting the metabolism of host steroids related to the development of the immune response to infection, although existing data only partially confirm this link. Support for this hypothesis can be found in the study by Varaksa et al.[23], which experimentally demonstrated that enzymes of Mtb, including CYP124, can metabolise human immune oxysterol messengers. This work, along with studies[22,77], also confirms CYP124 interaction with components of the vitamin D pathway (7-dehydrocholesterol, vitamin D3), whose role in antimicrobial immunity, particularly against Mtb, is covered in reviews[22,78]. Compounds such as 24-hydroxycholesterol [80,81]and (*S*)-2,3-epoxysqualene [79] are associated with immune or inflammatory processes, but their potential interaction with CYP124 has not yet been studied. Also, a number of publications [84,85,92] indicate a significant immunomodulatory role of sex hormones (including progesterone, DHEA, testosterone, pregnenolone, and estrogens) in infections; however, no studies were found on their binding to the CYP124 active site. The assumption that these steroid compounds, which lack a pronounced aliphatic chain at the C17 position, could be substrates of CYP124, must be treated very cautiously, considering the known specificity of CYP124 for preferential ω-hydroxylation of methyl-branched lipids[20]. Thus, despite the established role of CYP124 in the metabolism of individual immune-relevant host sterols, the hypothesis of its broader involvement in immune regulation requires further verification.

Thus, we consider our selection of priority 9 compounds as examples of chemical structures that may be characteristic of both substrates from classical human endogenous metabolites and prototypes of CYP124 inhibitors. Based on this, analysis of the ligand-CYP124 complex structure appears interesting. This can be useful as a source of information on the involvement of amino acids lining the walls of the CYP124 active site. This information may be important in the search for inhibitor prototypes, facilitating the determination of compound modification pathways [20]. To predict the structure of complexes of the 9 ligands with CYP124, we used soft docking in the Flare program.

As a result, 8 models of compound-CYP124 complexes were obtained for four compounds (medicarpin, maackiain, 3,26-di-OSO_3_-octaol, 3-OSO_3_-octaol) in three variants of CYP124 crystal structures (Table 3). For the other 5 compounds, no acceptable docking poses were identified in any of the CYP124 crystal structures. Different CYP124 crystal structures were used due to the significant conformational mobility of the active site cavity, the configuration of which is significantly influenced by the structure of the bound compound [22,23,77].

The low proportion of successful docking results can be explained by the conformational mobility of the CYP124 active site: soft docking does not account for significant structural shifts in the protein molecule. Due to these circumstances, MD of HD-4 and 15-β octaol complexes with Mtb CYP124 was performed. Computational results demonstrate tighter heme proximity and more favourable interactions for HD-4 versus 15β-octaol, aligning with spectral titration data. For 15β-octaol, noncovalent interactions anchor the ligand but likely minimally impact spectral alterations. Instead, spectroscopic properties could appear to be governed by hydrophobic/H-π interactions of the C25/26-OH groups with heme and the water-mediated 26-OH–Thr271 H-bond. Collectively, these results elucidate possible CYP124’s ligand-specific interaction mechanisms, highlighting distinct binding modalities between sterol derivatives.

MD revealed a gain in energy upon the complex formation with HD-4 and 15β-octaol of -32.85 kcal/mol and -10.10 kcal/mol, respectively. Despite this significant (approximately threefold) difference in binding energies calculated by MD simulation, the experimental dissociation constants (Kd) obtained from spectral titration were relatively close: 4.06 µM for HD-4 and 10.37 µM for 15β-octaol. To compare these results directly, we estimated the energy change during the CYP124 interaction with HD-4 and 15β-octaol during the spectral titration via Van ’t Hoff equation ΔG = RT ln Kd, using the T = 298 K (25 °C), R = 8.31 J⋅K^−1^ mol^−1^. For HD-4 ΔG was -5.98 kcal/mol, for 15β-octaol – -5.43 kcal/mol. Both experimental data and computational modeling demonstrate a unidirectional change in Gibbs free energy for the interaction of both compounds and indicate that HD-4 exhibits stronger binding to the protein compared to 15β-octaol. The discrepancy between the experimental and calculated Gibbs free energy changes (approximately 5-fold) for HD-4 may be attributed to the presence of a carbohydrate substituent, which could bias the results of molecular modeling [113,114].

Next, the list of CYP124 active site amino acid residues interacting with low molecular weight compounds, both in our obtained models and in crystal structures, was analysed. Such analysis is important for understanding the mechanisms of substrate binding in the CYP124 active site, as well as for optimising pathways for chemical modification of potential inhibitor chemical structures.

Comparison of amino acid interactions with CYP124 ligands, according to crystallography data (PDB ID: 6T0F, 6T0G, 6T0K), molecular docking, and MD, revealed a conserved hydrophobic core, including 11 residues (Phe63, Thr95, Leu103, Phe107, Ile111, Leu198, Phe212, Val266, Ala267, Met318, Phe416). The critical role of Phe107 (π-stacking), Phe416 (hydrophobic pocket), Val266/Ala267 (“hydrophobic wall”), and Ile111/Met318 (stabilisation of substrate binding to heme) is consistent with literature data on substrate binding by CYP124 *M. tuberculosis*[20].

Specifically, Johnston et al. showed that 7 residues (Ile94, Leu103, Phe107, Ile111, Leu263, Val266, Ala267) form a universal core, positioning the terminal methyl groups of lipids at a distance of 3.8 Å from the heme for ω-hydroxylation[20].

Consistently in docking and dynamics, but not in crystals, Ile94, Leu263, Thr271, Asn93, Asn97, Ser109 appear, forming an adaptive hydrophobic pocket (Ile94, Leu263) and an H-bond network (Thr271, Asn93, Asn97, Ser109) for orienting polar ligand groups. The role of Ile94, Leu263, and Asn97 in forming hydrophobic interactions and hydrogen bonds, respectively, with CYP124 ligands is supported by literature data[20].

Unique MD residues (Glu270, Phe112, Phe200, Phe209, Phe216) partially overlap with amino acids responsible for substrate-induced closure of the enzyme active site (Phe200, Phe212) according to crystallography data[20], explaining their incomplete reproduction in docking due to static modelling limitations. Dynamics of Thr100–Phe107 (BC-loop) and Phe209–Val231 (G-helix) in MD corresponds to conformational reorganisation upon binding [20]. Polar residues Thr95 and Asn97 exhibit adaptability in MD: Thr95 participates in hydrophobic contacts (-2.47 kcal/mol) and H-bonds with the 6β-hydroxy group of HD-4, and Asn97 forms H-bonds (-2.3 kcal/mol) with the 4β-OH of 15β-octaol, consistent with their role in stabilising polar substrate groups in crystals[20]. Unique residues found exclusively in molecular docking include Ile262, Ser259, Ser414, Thr100, Tyr106, Val112, Val315, Val64, participating in hydrophobic contacts and H-bonds, but absent in crystals and MD.

The absence of unique residues in the crystal structures used for modelling confirms their representativeness as a reference, and the discrepancies most likely reflect methodological features: docking captures local interactions, MD captures the dynamics of substrate-induced site closure, and their overlap reveals hidden aspects of binding. Moreover, these discrepancies do not contradict the dominant role of the enzyme’s active site hydrophobic pocket in protein-ligand interactions.

Thus, it can be stated that the conserved hydrophobic core of CYP124 (including Phe107, Phe416, Val266/Ala267) is the basis for ligand binding, as confirmed by crystallography, docking, and MD data, as well as literature. The present study, based on a comprehensive experimental and computational approach, yielded a number of significant results concerning the interaction of natural compounds and potential endogenous ligands with CYP124. These results form the basis for the following conclusions.

## 5. CONCLUSIONS

We analysed a sample of 32 compounds isolated from representatives of the flora and fauna of the Far East. Comprehensive analysis using surface plasmon resonance, spectral titration, and biochemical analysis allowed the identification of 9 new non-azole ligands for the active site of CYP124 *M. tuberculosis.* Among them, 15β-octaol (IC50 ≈ 86 μM) and HD-4 (IC50 >100 μM) acted as inhibitors forming long-lived complexes (τ1/2 of the CYP124/inhibitor complexes was 181 and 65 min, respectively). The spectrum of biological activities was predicted for all detected enzyme active site ligands, revealing potential toxic risks (immunosuppression, cytotoxicity), which must be considered in case of further optimisation of these structures. Thus, a foundation was laid for the search for new non-azole inhibitors of Mtb CYP124 and natural compounds were identified that can serve as base structures for creating modifiers of the enzymatic activity of the cytochrome.

By analysing the KEGG database, which is a collection of small molecules, biopolymers, and other chemical substances relevant to biological systems, 37 human endogenous metabolites structurally similar to the found natural CYP124 ligands (Tanimoto coefficient >0.4) were discovered, including immunoregulatory sterols and their precursors, among which some are already known as CYP124 active site ligands (7-dehydrocholesterol, vitamin D3, cholesterol and 25-hydroxycholesterol), consistent with previously known data on the enzyme’s possible role in host steroid metabolism during infection. Presumably, such immunoregulatory compounds and their precursors: previtamin D3, calcidiol, calcitriol, 24-hydroxycholesterol and (S)-2,3-epoxysqualene may also be ligands of the CYP124 active site, but this assumption requires experimental confirmation.

*In silico*-analysis (docking, molecular dynamics) and its comparison with known CYP124 crystal structures confirmed the role of the CYP124 hydrophobic core (Phe63, Thr95, Leu103, Phe107, Ile111, Leu198, Phe212, Val266, Ala267, Met318, Phe416) in ligand binding. At the same time, unique residues (Glu270, Phe112, Phe200, Phe209, Phe216) not participating in ligand interaction in the investigated crystal structures were revealed in molecular dynamics models. This discrepancy likely reflects substrate-induced conformational changes and requires further study.

The obtained data on binding patterns create a basis for pharmacophore modelling in the development of inhibitors of one of the potentially key enzymes of *M. tuberculosis*, which may prove to be a new approach in the fight against multidrug-resistant tuberculosis.

## 6. AKNOWLEDGEMENTS

We thank Prof. Rita Bernhardt (Saarland University, Saarbrucken, Germany) for providing the expression construct for Arh1.

Surface plasmon resonance analysis was performed using the equipment of the Human Proteome Core Facilities of the V.N. Orekhovich Research Institute of Biomedical Chemistry (Moscow, Russia).

## 8. AUTHOR CONTRIBUTIONS

Conceptualization – LK and EY, Methodology – LK, EY, TV, AG, AK, IG, YM, EZ, OG, DT, TM, SK, AK, EV, NM, AS, SA, TR and DT; Data curation and Formal analysis – LK, EY and YM; Visualization – LK, EY, TV and EZ; Writing - original draft – EY; Writing - review & editing – LK, EY and YM; Supervision – VK, AK, EK, SF, NI, PD, AG and AI; Funding acquisition – PD and AG; All authors have read and agreed to the published version of the manuscript.

