## Supplementary data for "Diverse Ligands for Mycobacterial CYP124 Identified from Plant and Marine Compounds"

### 1. The detailing of the compounds purification protocols.

The studied compounds were isolated or obtained by chemical modifications at the G.B. Elyakov Pacific Institute of Bioorganic Chemistry, Far Eastern Branch of the Russian Academy of Sciences, similar to the protocols published earlier [1–13] from the starfish *Henricia derjugini*, *Henricia leviuscula spiculifera*, *Archaster typicus*, *Lethasterias nanimensis chelifera*, and *Patiria pectinifera*, the holothurian *Colochirus robustus*, the sea sponge *Rhabdastrella globostellata*, the crinoid *Phanogenia gracilis*, the sea urchins *Scaphechinus mirabilis* and *Mesocentrotus nudus*, the Far Eastern endemic plant *Maackia amurensis*, and the sea grass of the Zosteraceae genus. The purity of the isolated compounds was confirmed by HPLC, NMR spectroscopy ($^1$H and $^{13}$C NMR) and mass spectrometry in comparison with the samples described in the cited works.

Thus, spiculiferosides A–C were isolated from the starfish *Henricia leviuscula spiculifera* according to the following original protocol [1]. Freshly collected specimens of starfish *H. leviuscula spiculifera* were immediately frozen after fishing. The sliced specimens (1.1 kg) were extracted twice with EtOH at room temperature. The extract was evaporated under reduced pressure and the residue (83.2 g) was dissolved in $H_2O$ (0.5 L). The $H_2O$-soluble fraction was passed through a Polychrome-1 column (7.5 × 75 cm) and eluted with $H_2O$ and then with EtOH. The combined EtOH eluate was concentrated under reduced pressure and the resulting total fraction (9.7 g) was chromatographed over a Si gel column (6.5 × 15 cm) using $CHCl_3$/EtOH (stepwise gradient, 5:1→1:3, v/v). The obtained fractions were further purified on Florisil columns (7 × 15 cm) using $CHCl_3$/EtOH (stepwise gradient, 3:1→1:3, v/v) to yield four main fractions 1–4. Fraction 2 (641 mg) was separated by HPLC on a Discovery C18 column (MeOH/$H_2O$/1M $NH_4OAc$, 55:44:1, v/v/v, flow rate 1.7 mL/min) to give pure spiculiferosides B and C. Fraction 3 (235 mg) were subjected to HPLC on an YMC-Pack Pro C18 column (54% aq. EtOH, flow rate 2.0 mL/min) and purified repeatedly on the same column (50% aq. EtOH, flow rate 2.0 mL/min) to afford pure spiculiferoside A. Steroidal compounds from the starfish *Henricia derjugini*, *Archaster typicus*, and *Patiria pectinifera* were isolated using similar protocols [2–4].

Asterone (Agl1) was isolated after hydrolysis of the total asterosaponin fraction using the following method [5]. The total asterosaponin fraction was isolated from the ethanol extract of the starfish *Lethasterias nanimensis chelifera* collected off Shiashkotan Island (Kuril Islands, Pacific coast) using column chromatography on Amberlite XAD-2, Sephadex LH-20 and Silica gel. Acid hydrolysis was performed with 2 N HCl upon heating at 100 °C for 2 hours. Aglycons were extracted with chloroform. The resulting total asterogenins were sequentially separated on Silica gel columns in the $CHCl_3$/EtOH (18:1) system and in the $CHCl_3$/EtOAc (30:1 x 30:15) system. Further separation was performed by HPLC on Zorbax ODS (5 μm, 250 x 4.6 mm) and Separon SGX (5 μm, 150 x 3.0 mm) columns. As a result, asterone (Agl1) was obtained as the main aglycon.

Sulfated steroids (25*S*)-5α-cholestane-3β,4β,6α,7α,8,15α,16β,26-octaol 3-*O*-sulfate (3-$OSO_3$-octaol) and (25*S*)-5α-cholestane-3β,4β,6α,7α,8,15α,16β,26-octaol 3,26-*O*-disulfate (3,26-di-$OSO_3$-octaol) were obtained by sulfation of (25*S*)-5α-cholestane-3β,4β,6α,7α,8,15α,16β,26-octaol (15α-octaol) isolated from the starfish *Patiria pectinifera* as described in the article [4]. Sulfation of (25*S*)-5α-cholestane-3β,4β,6α,7α,8,15α,16β,26-octaol was carried out with pyridine sulfotrioxide ($C_5H_5N{\cdot}SO3$) in dry pyridine. For more complete sulfation of the initial steroid compound, a slight excess (by weight) of pyridine sulfotrioxide was used. The reaction was carried out at 60 °C for 24 hours. 500 μL of 10% KOH were added to the cooled reaction mixture to neutralize the solution. After neutralization, the resulting mixture of derivatives was purified from salts on a column with Polychrome-1 (1.5 × 10 cm). The adsorbed substances were washed off with pure ethanol and evaporated *in vacuo*. The resulting mixture of derivatives was separated by HPLC on a Discovery HS C18 reversed-phase column in a 70% MeOH system at a flow rate of 2.5 mL/min. The fraction was dissolved in methanol. As a result, (25*S*)-5α-cholestane-3β,4β,6α,7α,8,15α,16β,26-octaol 3-*O*-sulfate (3-$OSO_3$-octaol) and (25*S*)-5α-cholestane-3β,4β,6α,7α,8,15α,16β,26-octaol 3,26-*O*-disulfate (3,26-di-$OSO_3$-octaol) were obtained.

The triterpene glycoside hemioedemoside A was isolated from the sea cucumber *Colochirus robustus* following the original protocol [6]. Samples of the sea cucumber *C. robustus* (family Cucumariidae; order Dendrochirotida) were collected from Nha Trang Bay, South China Sea. The sea cucumbers were ground and extracted twice with boiling 60% ethanol. The dry residue weight was about 23.5 g. The ethanol extract of *C. robustus* concentrated *in vacuo* was chromatographed on a Polychrome-1 column, eluting first the inorganic salts and impurities with $H_2O$, and then the glycosides with 50% ethanol. Then chromatography was performed on Silica gel columns using the $CHCl_3$/EtOH/$H_2O$ system (100:125:25 and 100:100:17) as mobile phases to obtain a number of glycoside subfractions. Their further separation was carried out by HPLC on a Supelco Ascentis RP-Amide semi-preparative reversed-phase column (10×250 mm, 5 µm), hemioedemozide A was isolated from subfraction 4 using the acetonitrile/$H_2O$/1M $NH_4OAc$ (38:60:2) solvent system.

The isomalabarican triterpenoids stellettins Q and R, and nor-isomalabaricanes jaspolide F and globostellettin G were isolated from the Vietnamese marine sponge *Rhabdastrella globostellata* as described in [7,8]. The sponge was assigned to the genus *Stelletta* and then re-identified and reported as *R. globostellata* species [14]. The EtOH extract of defrosted and chopped sponge was concentrated and partitioned between $H_2O$ and EtOAc. The organic layers were separated using LH-20 ($CHCl_3$/EtOH, 1:1) and silica gel (step-wise gradient $CHCl_3$→EtOH) columns to give nine fractions. Fourth fraction contained isomalabaricanes that were purified by column re-chromatography (silica gel and YMC-Pack ODS-A) followed by HPLC. Final isolation procedures for the light-sensitive isomalabaricanes were performed using amber glassware and aluminum foil wrapping to avoid interconversion of *Z*/*E*-isomers. Stellettins Q and R were obtained by the reversed-phase HPLC (Discovery HS F5-5, 10 × 250 mm, 5 µm) in 70% and 80% EtOH correspondingly. To purify *Z*/*E*-isomers jaspolide F, and globostelletin G we used normal-phase HPLC (Ultrasphere-Si, 10 mm × 250 mm, 5 µm, $CHCl_3$/EtOH, 35:1)

A quinoid compound, phanogracilin A (570-2), was isolated from the crinoid *Phanogenia gracilis* according to the original protocol [9]. *P. gracilis* was collected from the South China Sea near Ly Son Island in May 2021. Samples (58.5 g) were extracted with 70% ethanol containing 10% $H_2SO_4$ to obtain a dark red solution with a green tint. The acidified EtOH extract was concentrated *in vacuo*, and the residue was then extracted with chloroform and ethyl acetate. HPLC-MS analysis showed that the chloroform and ethyl acetate extracts contained the same pigments in slightly different ratios, which were present in trace amounts in the ethanol extract: two compounds with molecular weights of 570 and one compound with a molecular weight of 542. The combined chloroform and ethyl acetate extracts (0.8 g), obtained from the acidified EtOH extract, were repeatedly chromatographed on a silica gel column, where the silica gel was pre-impregnated with a few drops of 5 mg/mL oxalic acid in ethanol. The column was eluted with hexane/$CHCl_3$ with gradually increasing amounts of chloroform (hexane/$CHCl_3$ 1:0, 3:1, 2:1, 1:1, 1:2, 1:3, 1:4, 1:5) and then with $CHCl_3$/MeOH with gradually increasing amounts of MeOH ($CHCl_3$/MeOH 100:1, 50:1, 50:3, 10:1, 9:1, 5:1, 1:1, 1:0). As a result, a new bibenzochromenone phanogracillin A was obtained.

A quinoid pigments echinochrome A, echinamines A and B were isolated from the sea urchin *Scaphechinus mirabilis* Agassiz, and spinochromes B, D, and E were obtained from the sea urchin *Mesocentrotus nudus* according to the original protocol [10,11]. The defrosted sea urchins *Scaphechinus mirabilis* (A. Agassiz, 1864) (2 kg, wet wt) were extracted with EtOH containing 10% $H_2SO_4$ (3 L) at room temperature. The EtOH extract were concentrated *in vacuo*. The residue was partitioned between $H_2O$ (0.3 L) and $CHCl_3$ (3×0.3 L). Quinoid pigments as sodium salts were extracted from $CHCl_3$ with 1% $Na_2CO_3$ solution (0.3 L, under argon). The solution of pigment sodium salts was acidified to pH 2, and quinones were extracted with $CHCl_3$ (2×0.2 L) and EtOAc (2×0.2 L). The combined organic extracts were evaporated, and the residue (406 mg) was sequentially separated by low-pressure on a Toyopearl HW-40 column (20×2 cm) using 20-50% EtOH containing 0.5% HCOOH with gradient elution to obtain three fractions. Fraction, containing echinochrome A was purified in the same manner, followed by column chromatography (50×1 cm) on a Sephadex LH-20 column, using 7:1 chloroform/ethanol, to give echinochrome A. Fraction 3 contained most of echinamines A and B as assessed by HPLC analysis. This fraction was separated by chromatograraphy on a Toyopearl HW-40 column (40×2 cm) using 40-60% EtOH containing 0.5% HCOOH, giving echinamine A and echinamine B. The acidified EtOH extract (2.0 L) of defrosted *Mesocentrotus nudus* shells and spines (1.8 kg) was concentrated *in vacuo* at 55 °C and partitioned between $H_2O$ and EtOAc. The EtOAc fraction was subjected to chromatography using a Toyopearl HW-40 column. The column was eluted with $H_2O$ – EtOH (containing 0.5% HCOOH), with gradually increasing amounts of EtOH. All the colored fractions were analyzed by HPLC-DAD-MS and chromatographed repeatedly under the same conditions on a YMC-Pack ODS-A column to yield spinochrome B, spinochrome D, and spinochrome E.

The isolation of polyphenolic compounds from *Maackia amurensis* was carried out according to the following method [12]. The *M. amurensis* heartwood (500 g) was extracted twice with a mixture of $CHCl_3$–EtOH at a ratio of 3:1 for 3 h (60 °C). The air-dried extract (15.5 g) was applied to a polyamide column (50–160 μm) and eluted with hexane/$CHCl_3$ and $CHCl_3$/EtOH solution systems to obtain fractions 1–8 and 9–18, respectively. We gradually increased the portions of $CHCl_3$ and EtOH in the solution systems (hexane/$CHCl_3$, *v/v*: 1:0, 10:0, 8:1, 5:1, 2:1, 1:1, 1:2, $CHCl_3$; $CHCl_3$/EtOH, *v/v*: 1:0, 100:1, 50:1, 40:1, 30:1, 20:1, 10:1, 5:1, 2:1). The fractions containing polyphenolic compounds according to HPLC data were selected for further purification. Fraction 9 (670 mg) was then chromatographed twice on a silica gel column (40–63 μm) to obtain pure compounds of retusin and maackiain. Fraction 10 (895 mg) was also chromatographed twice on a silica gel column to obtain the individual compounds of tectorigenin, medicarpin, formononetin, and (±)-3- hydroxyvestitone. Liquiritigenin was isolated from fraction 11 (610 mg) by silica gel chromatography. Fraction 13 (740 mg) contained the compounds maackin and piceatannol. These polyphenols were separated on a silica gel column and then purified on a C-18 column (YMC gel ODS-A 75 μm).

Luteolin 7,3′-disulfate was obtained from marine plants of the genus Zosteraceae as described in [13]. Freshly harvested green eelgrass (Zostera spp., family Zosteraceae) were desalinated using potable water. Mechanical impurities, algae, and other marine plant species were removed. The Zostera was washed three times and then placed on a mesh filter until the water drained completely. The drained Zostera was loaded into a reactor under weights to prevent flotation and covered below the meniscus with 96% ethanol (raw material : extractant ratio 1:(1-2)). Extraction was carried out for 12-24 hours.

The ethanolic extract was decanted and filtered through fabric, paper, or cotton wool filter media. This extraction process was repeated three times. The combined ethanolic extracts were concentrated under reduced pressure. The resulting concentrate was dissolved in water. The solution was centrifuged or filtered, and the sediment was discarded. The filtrate was acidified with 15-20% hydrochloric acid to pH 1-2 and left for 24 hours at 2-4 °C to allow formation of the acid-insoluble lignin precipitate. The precipitate was separated by centrifugation or filtration. The acidic solution of phenolic compounds was applied to a column packed with Polychrome-1, pre-equilibrated with distilled water. Polyphenolic compounds bound to the Polychrome-1. Mineral salts and hydrochloric acid were removed by washing the column with distilled water. Elution of the bound polyphenolic compounds was performed using an ethanol gradient. The most polar polyphenol, luteolin 7,3'-disulfate, was eluted with 5% aqueous ethanol solution. The aqueous-ethanolic eluate was concentrated under reduced pressure at 60 °C until ethanol was completely removed. The aqueous residue was dried by either freeze-drying (lyophilisation) or spray-drying to constant weight.

**Table S1.** Difference spectra of CYP124 in the presence of compounds tested.

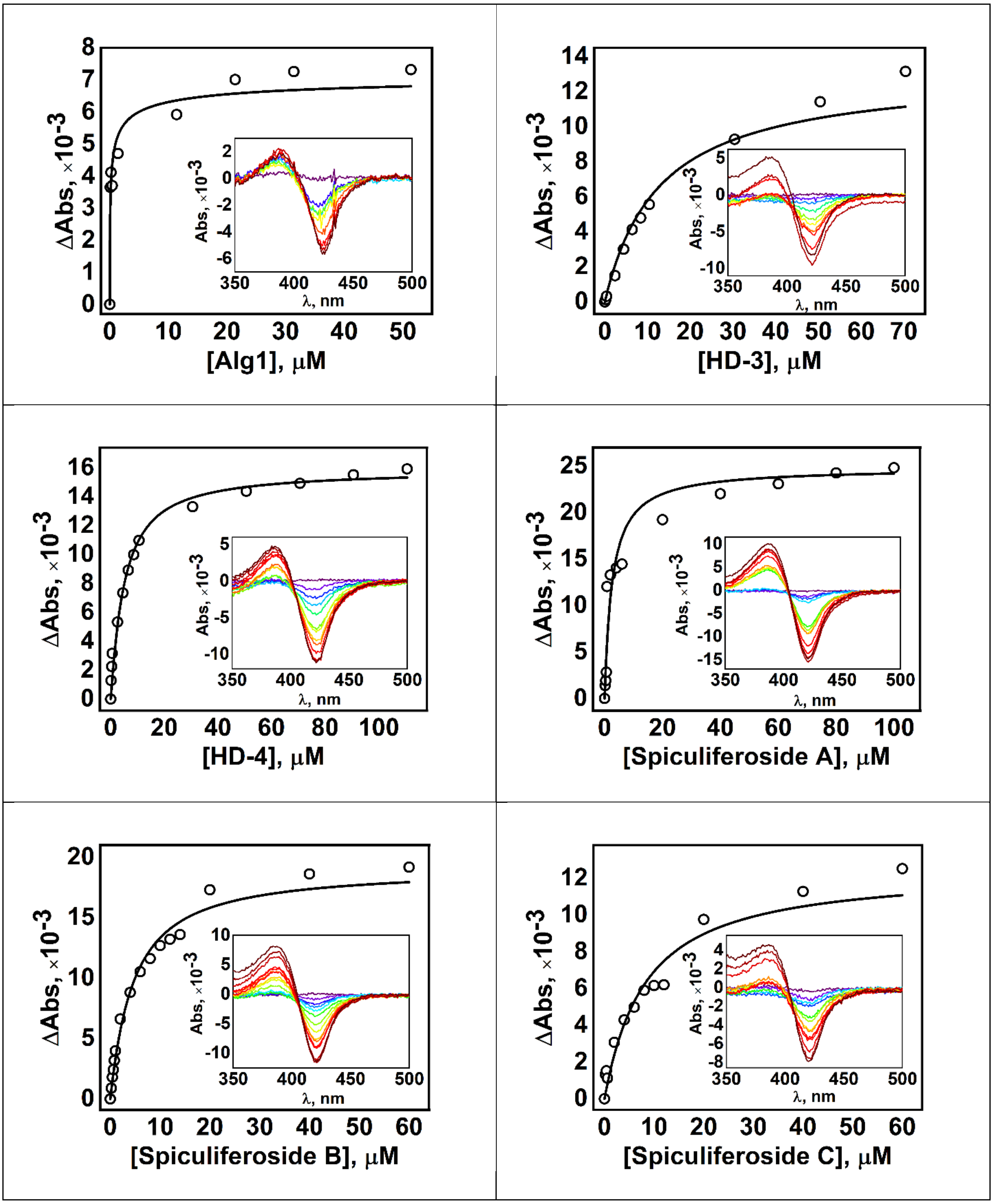

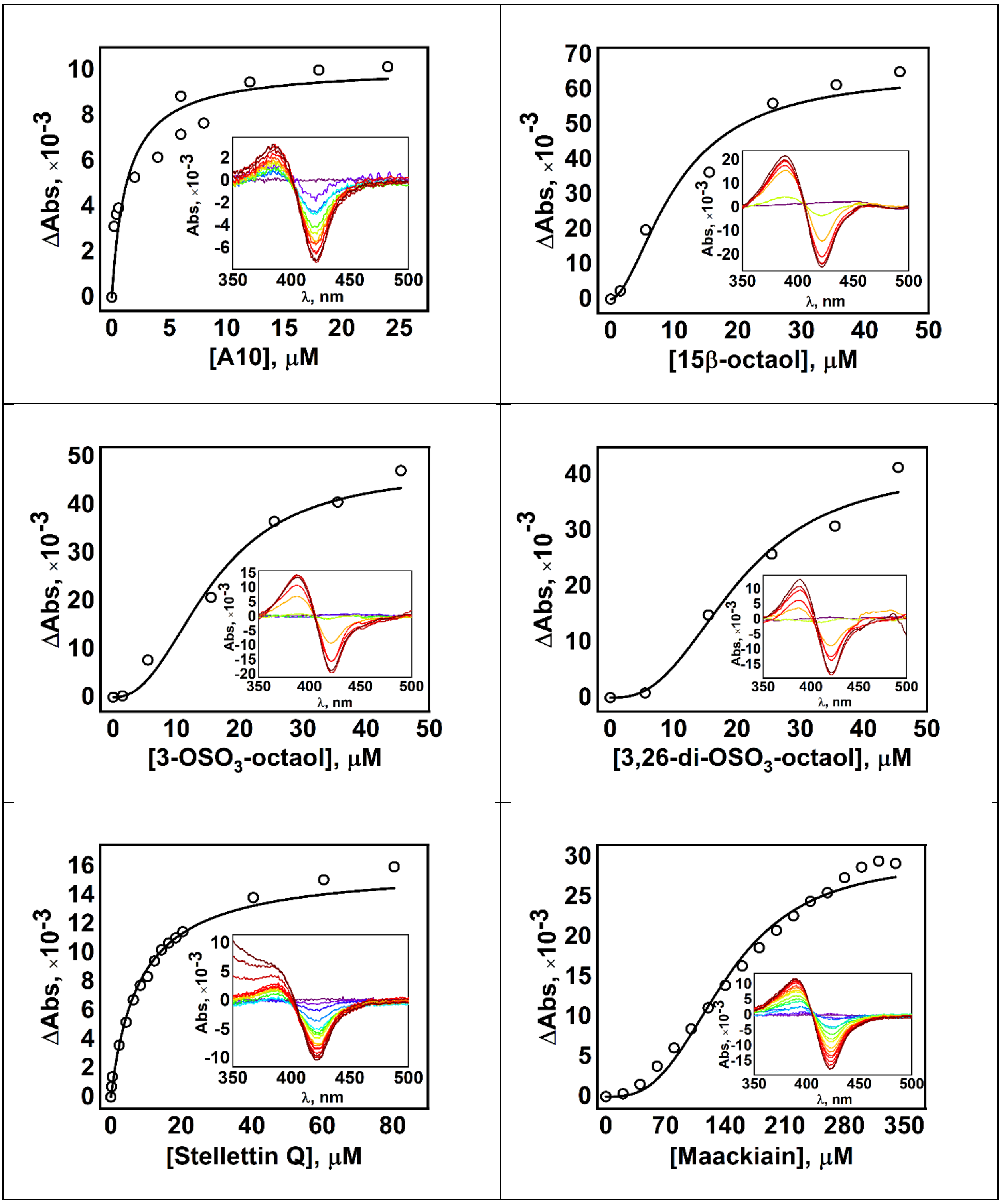
ΔAbs, ×10-3
[A10], μM
Abs, ×10-3
λ, nm
[15β-octaol], μM
[3-OSO3-octaol], μM
[3,26-di-OSO3-octaol], μM
[Stellettin Q], μM
[Maackiain], μM

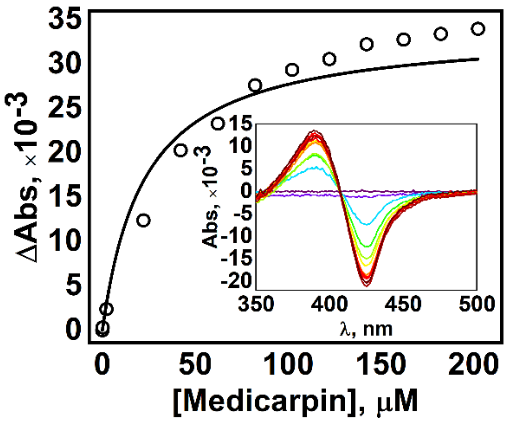

ΔAbs, ×10-3
35
30
25
20
15
10
5
0
0
50
100
150
200
[Medicarpin], μM
Abs, ×10-3
15
10
5
0
-5
-10
-15
-20
350
400
450
500
λ, nm

**Table S2.** Prediction of biological activity spectra of hit compounds

| Pa | Pi | Activity name |
|---|---|---|
| *3.26-di-$OSO_3$-octaol* | | |
| 0.928 | 0.004 | Benzoate-CoA ligase inhibitor |
| 0.84 | 0.006 | Alkylacetylglycerophosphatase inhibitor |
| 0.785 | 0.001 | Endoglycosylceramidase inhibitor |
| 0.77 | 0.002 | <u>Coenzyme-B sulfoethylthiotransferase inhibitor</u> |
| *3-$OSO_3$-octaol* | | |
| 0.833 | 0.007 | Alkylacetylglycerophosphatase inhibitor |
| 0.791 | 0.018 | Benzoate-CoA ligase inhibitor |
| 0.705 | 0.007 | Adenomatous polyposis treatment |
| *Spiculiferoside B* | | |
| 0.866 | 0.009 | Benzoate-CoA ligase inhibitor |
| 0.806 | 0.005 | **Glyceryl-ether monooxygenase inhibitor** |
| 0.813 | 0.012 | Mannotetraose 2-alpha-N-acetylglucosaminyltransferase inhibitor |
| 0.787 | 0.004 | Hepatoprotectant |
| 0.803 | 0.029 | CDP-glycerol glycerophosphotransferase inhibitor |
| 0.736 | 0.004 | Mycothiol-S-conjugate amidase inhibitor |
| 0.708 | 0.006 | Chemopreventive |
| 0.712 | 0.015 | **Immunosuppressant** |
| 0.702 | 0.01 | <u>Antifungal</u> |
| *Spiculiferoside C* | | |
| 0.899 | 0.002 | Hepatoprotectant |
| 0.871 | 0.003 | **Glyceryl-ether monooxygenase inhibitor** |
| 0.873 | 0.008 | Benzoate-CoA ligase inhibitor |
| 0.842 | 0.004 | <u>Cholesterol antagonist</u> |
| 0.821 | 0.011 | Mannotetraose 2-alpha-N-acetylglucosaminyltransferase inhibitor |
| 0.781 | 0.004 | Chemopreventive |
| 0.803 | 0.029 | CDP-glycerol glycerophosphotransferase inhibitor |

| | | |
|---|---|---|
| 0.771 | 0.016 | Beta-adrenergic receptor kinase inhibitor |
| 0.771 | 0.016 | G-protein-coupled receptor kinase inhibitor |
| 0.755 | 0.01 | **Immunosuppressant** |
| 0.741 | 0.004 | Mycothiol-S-conjugate amidase inhibitor |
| 0.758 | 0.024 | Alkenylglycerophosphocholine hydrolase inhibitor |
| 0.722 | 0.002 | Endoglycosylceramidase inhibitor |
| 0.71 | 0.004 | **Bilirubin oxidase inhibitor** |
| | *HD-4* | |
| 0.871 | 0.005 | **Antineoplastic** |
| 0.864 | 0.008 | CYP3A4 substrate |
| 0.857 | 0.009 | CYP3A substrate |
| 0.821 | 0.004 | **Immunosuppressant** |
| 0.815 | 0.002 | Alcohol O-acetyltransferase inhibitor |
| 0.776 | 0.035 | CDP-glycerol glycerophosphotransferase inhibitor |
| 0.714 | 0.006 | Chemopreventive |
| | *Medicarpin* | |
| 0.911 | 0.003 | **Caspase 3 stimulant** |
| 0.898 | 0.001 | Chalcone isomerase inhibitor |
| 0.871 | 0.019 | Membrane integrity agonist |
| 0.859 | 0.018 | Aspulvinone dimethylallyltransferase inhibitor |
| 0.846 | 0.025 | CYP2C12 substrate |
| 0.818 | 0.017 | Chlordecone reductase inhibitor |
| 0.778 | 0.009 | Apoptosis agonist |
| 0.774 | 0.014 | TP53 expression enhancer |
| 0.769 | 0.016 | **Antineoplastic** |
| 0.754 | 0.002 | Skin whitener |
| 0.713 | 0.005 | Antiviral (Influenza) |
| 0.702 | 0.001 | Melanin inhibitor |
| 0.7 | 0.003 | NOS2 expression inhibitor |
| 0.71 | 0.016 | **JAK2 expression inhibitor** |
| 0.723 | 0.033 | Antiseborrheic |

| | | |
|---|---|---|
| *Maackiain* | | |
| 0.934 | 0.003 | **Caspase 3 stimulant** |
| 0.935 | 0.005 | Membrane integrity agonist |
| 0.788 | 0.013 | **Antineoplastic** |
| 0.751 | 0.011 | Apoptosis agonist |
| 0.722 | 0.004 | Antiviral (Influenza) |
| 0.722 | 0.022 | TP53 expression enhancer |
| *Stellettin Q* | | |
| 0.782 | 0.014 | **Antineoplastic** |
| 0.733 | 0.042 | Mucomembranous protector |
| *15β-octaol* | | |
| 0.895 | 0.005 | Alkenylglycerophosphocholine hydrolase inhibitor |
| 0.876 | 0.004 | Alkylacetylglycerophosphatase inhibitor |
| 0.853 | 0.008 | Acylcarnitine hydrolase inhibitor |
| 0.776 | 0.008 | Immunosuppressant |
| 0.782 | 0.014 | **Antineoplastic** |
| 0.747 | 0.008 | **Glyceryl-ether monooxygenase inhibitor** |
| 0.765 | 0.027 | Antieczematic |
| 0.748 | 0.013 | **Prostaglandin-E2 9-reductase inhibitor** |
| 0.727 | 0.003 | **DELTA14-sterol reductase inhibitor** |
| 0.716 | 0.006 | Adenomatous polyposis treatment |
| 0.711 | 0.051 | **Testosterone 17beta-dehydrogenase (NADP+) inhibitor** |

Pa – probability "to be active", Pi – probability "to be inactive". The most potentially hazardous biological activities and those not repeated for other studied compounds are highlighted in bold and underlined, respectively.

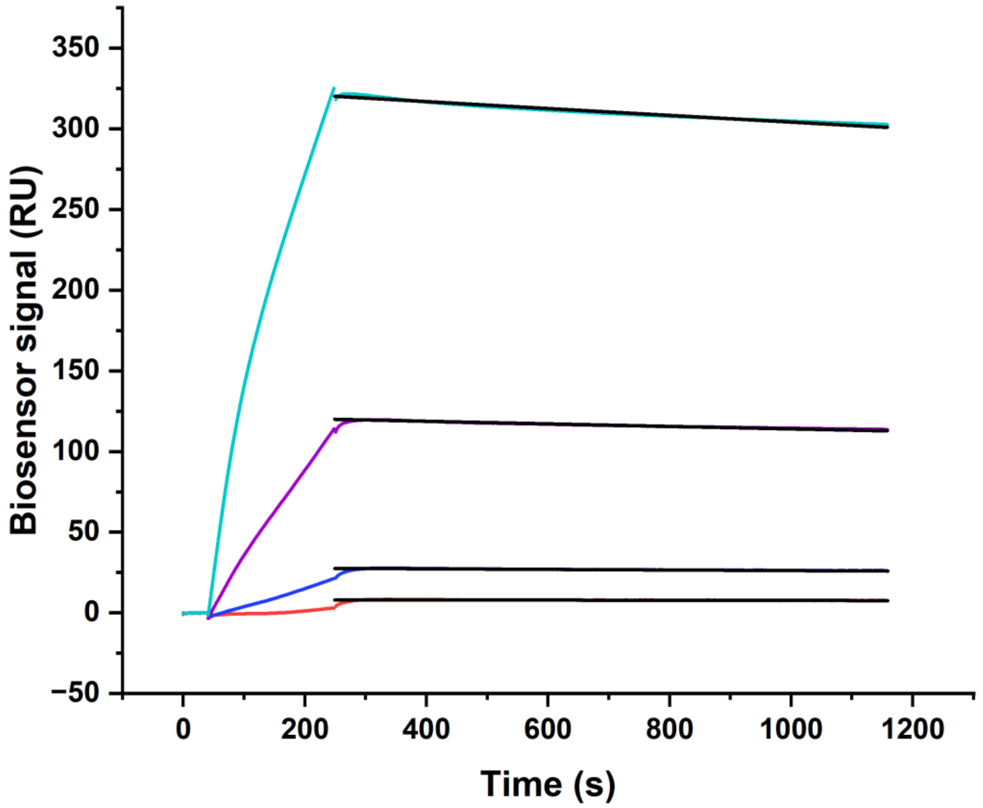


**Figure S1.** Typical surface plasmon resonance sensorgram of binding between immobilized CYP124 on the optical chip and 15β-octaol at different concentrations: 10 (red), 25 (blue), 50 (magenta), 75 (cian). Fitting curves (theoretical models) are highlighted in black.

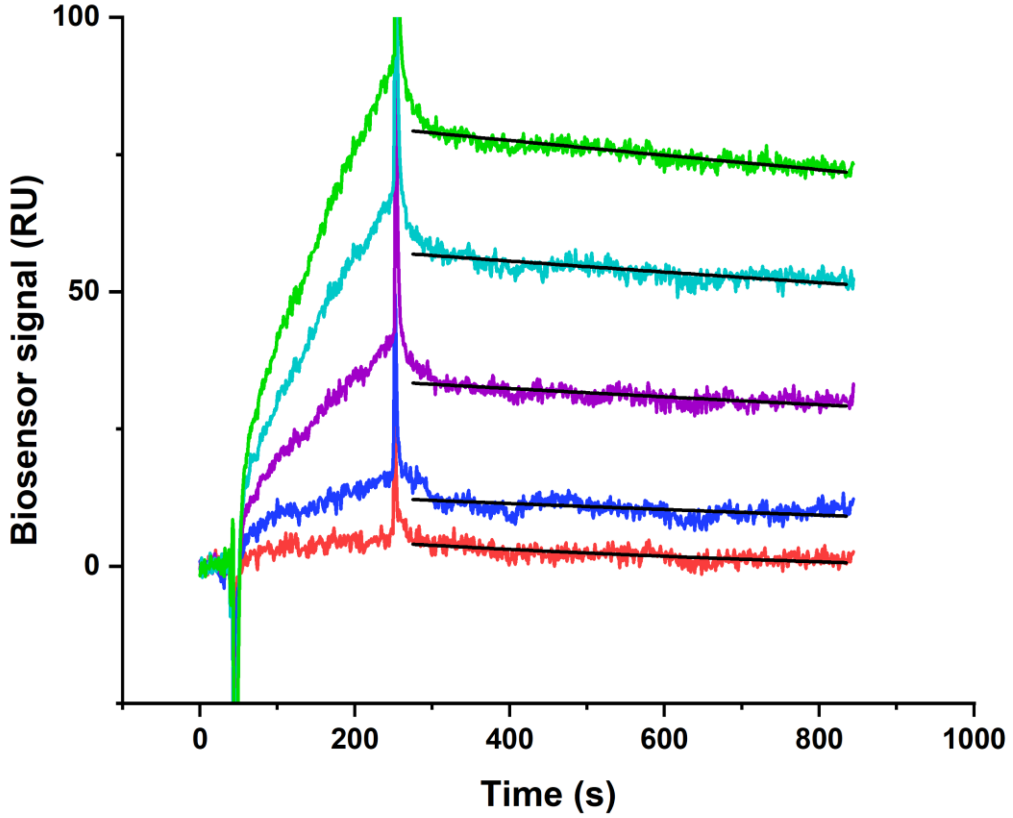


**Figure S2.** Typical surface plasmon resonance sensorgram of binding between immobilized CYP124 on the optical chip and HD-4 at different concentrations: 10 (red), 25 (blue), 50 (magenta), 75 (cian) and 100 µM (green). Fitting curves (theoretical models) are highlighted in black.

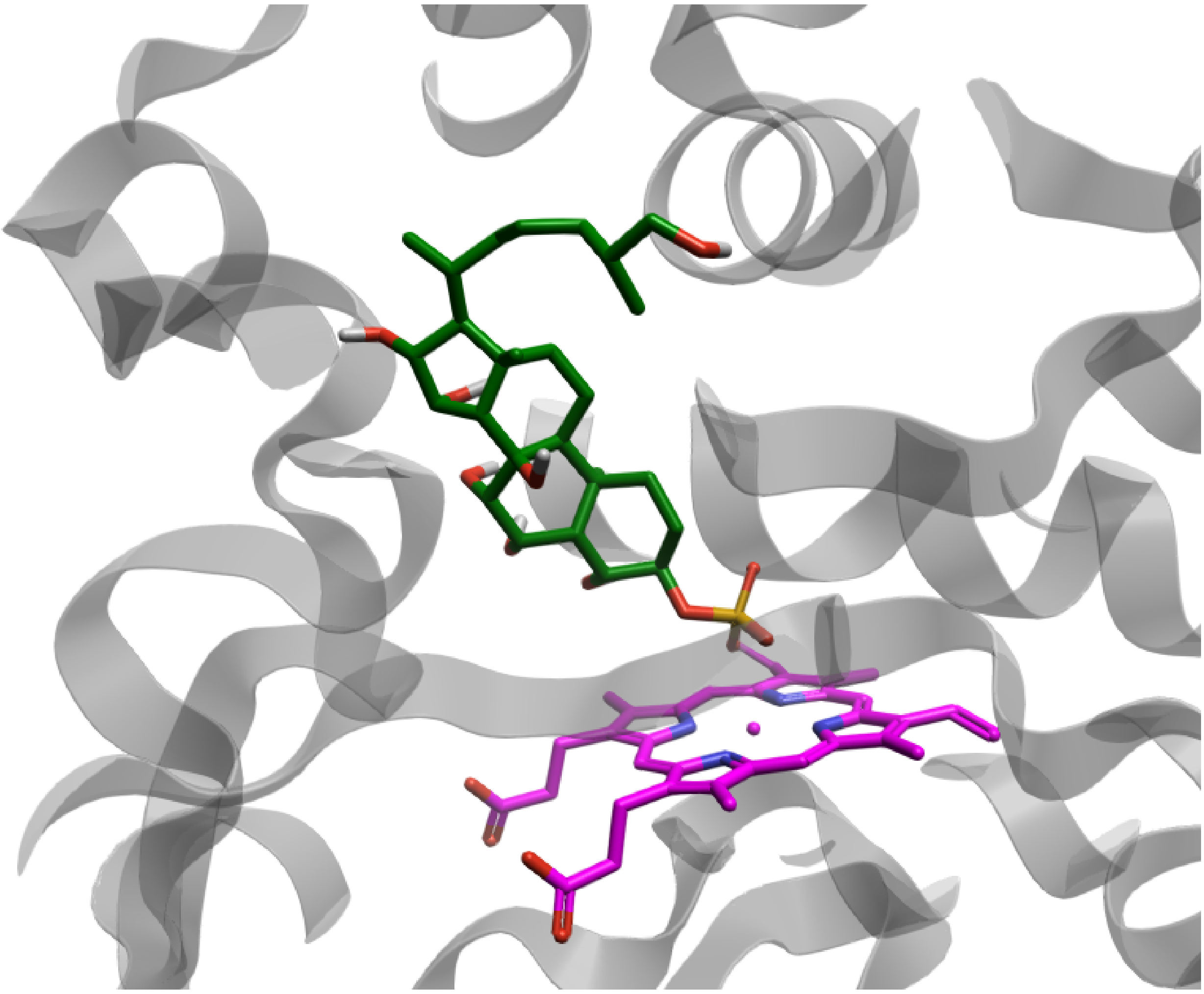

**Figure S3.** Diagram of the probable complex structures formed by CYP124 (structure's PDB ID 6T0K) with 3-$OSO_3$-octaol. The protein is depicted as a cartoon (grey) with the ligand represented as sticks (carbon – green, oxygen – red, phosphorus – yellow, hydroxyl's hydrogen – white), with heme shown in stick representation (carbon and ion – magenta, oxygen – red, nitrogen – blue).

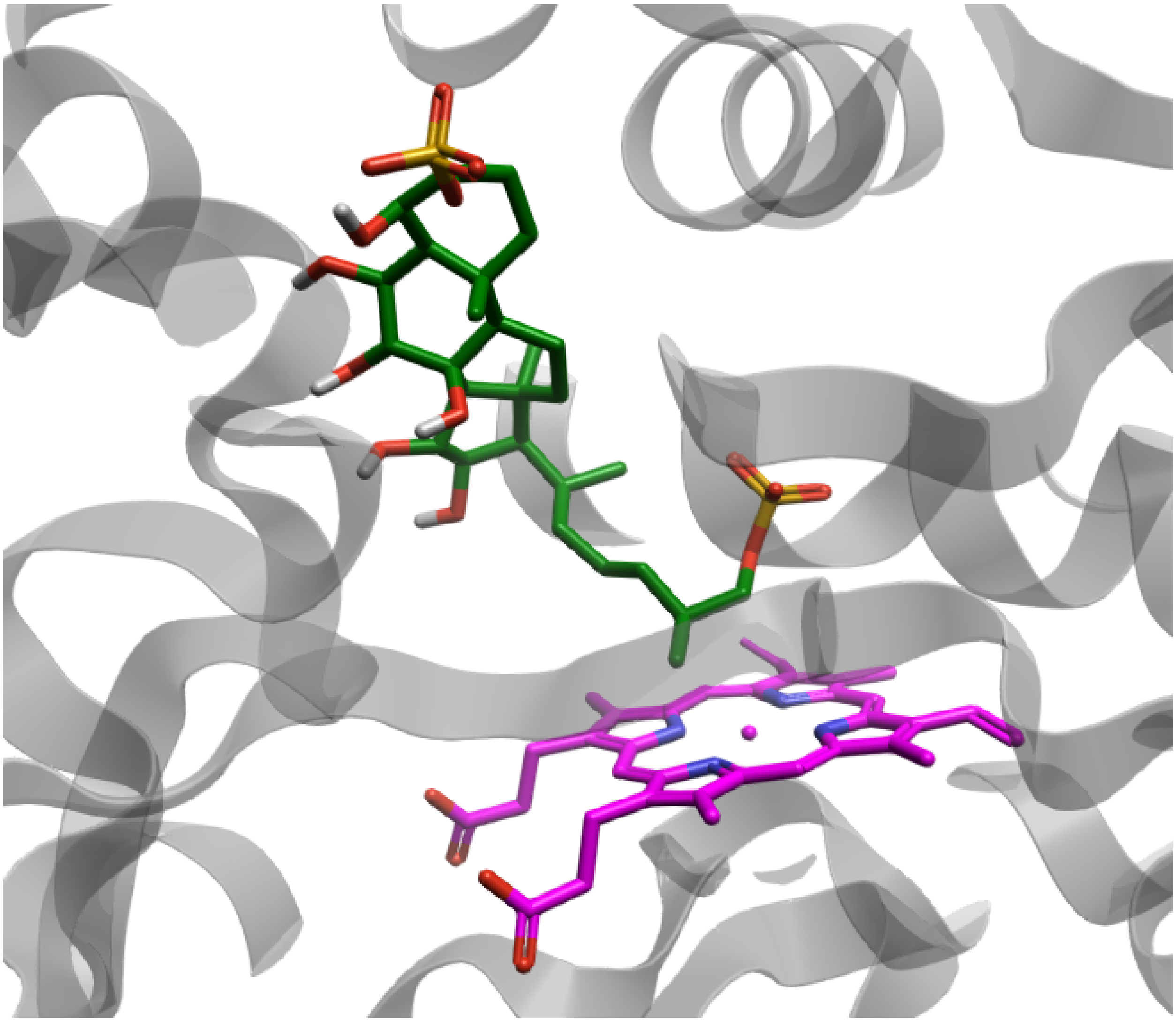

**Figure S4.** Diagram of the probable complex structures formed by CYP124 (structure's PDB ID 6T0K) with 3,26-di-$OSO_3$-octaol. The protein is depicted as a cartoon (grey) with the ligand represented as sticks (carbon – green, oxygen – red, phosphorus – yellow, hydroxyl's hydrogen – white), with heme shown in stick representation (carbon and ion – magenta, oxygen – red, nitrogen – blue).

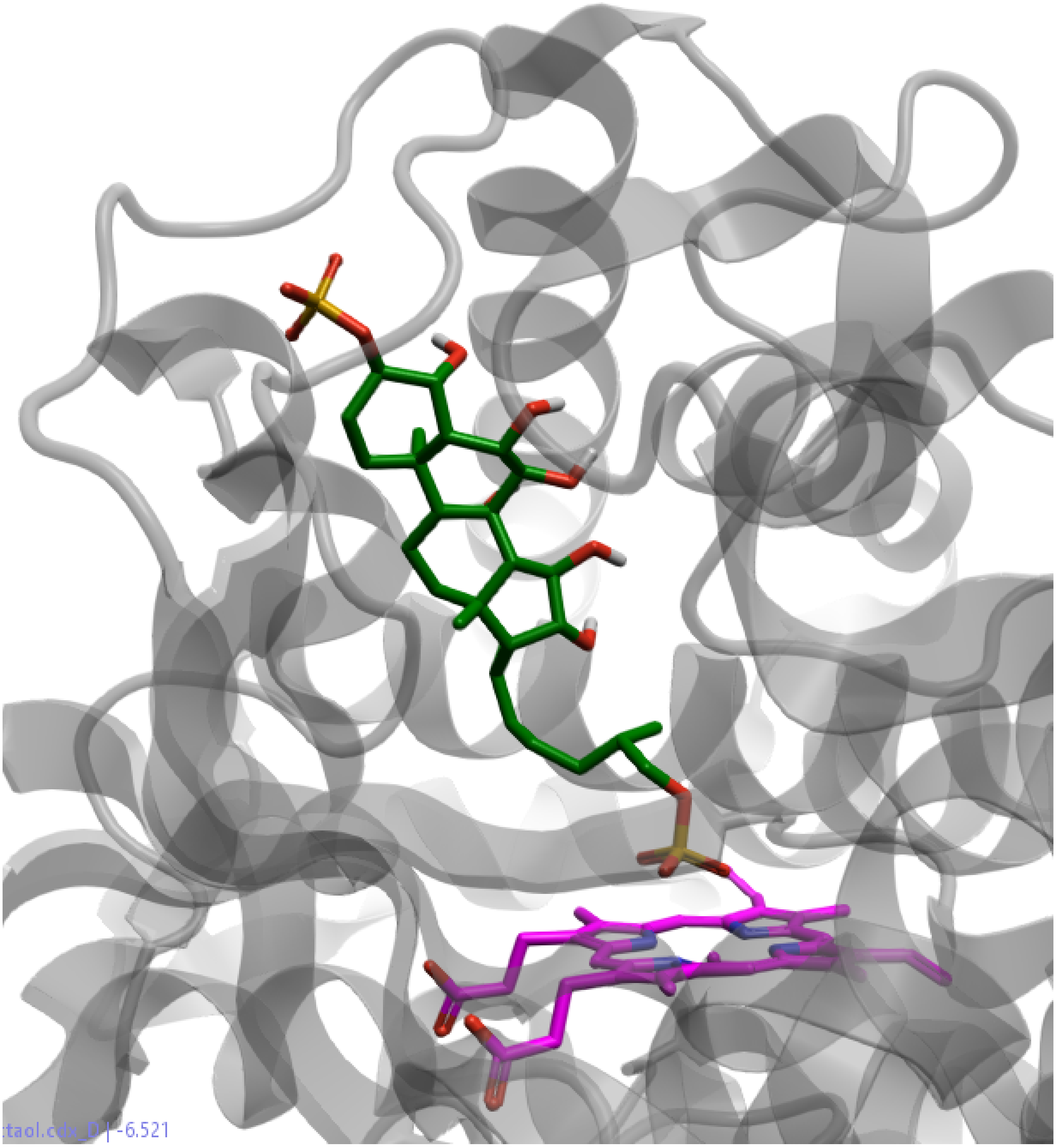


**Figure S5.** Diagram of the probable complex structures formed by CYP124 (structure's PDB ID 6T0F) with 3,26-di-$OSO_3$-octaol. The protein is depicted as a cartoon (grey) with the ligand represented as sticks (carbon – green, oxygen – red, phosphorus – yellow, hydroxyl's hydrogen – white), with heme shown in stick representation (carbon and ion – magenta, oxygen – red, nitrogen – blue).

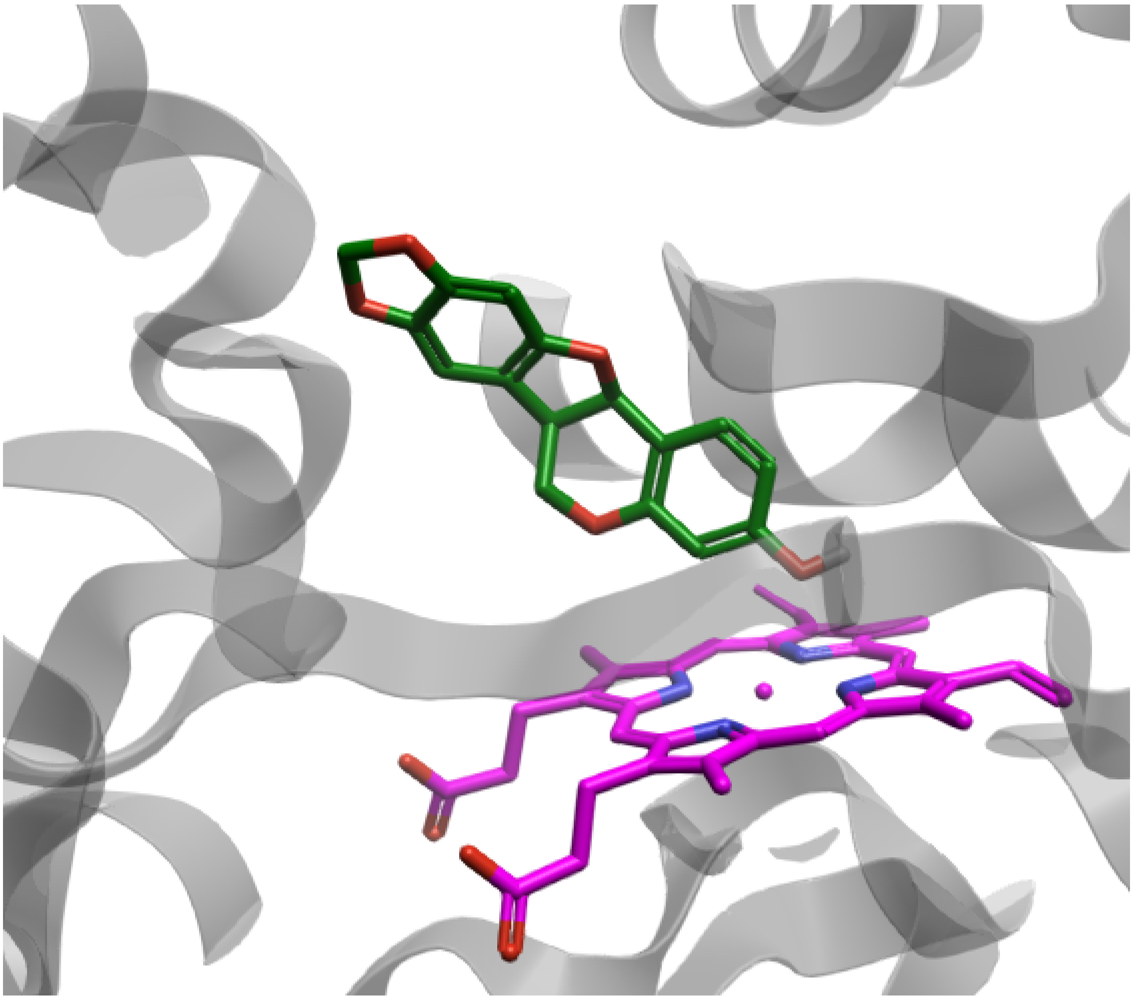

**Figure S6.** Diagram of the probable complex structures formed by CYP124 (structure's PDB ID 6T0K) with Maackiain. The protein is depicted as a cartoon (grey) with the ligand represented as sticks (carbon – green, oxygen – red, hydroxyl's hydrogen – white), with heme shown in stick representation (carbon and ion – magenta, oxygen – red, nitrogen – blue).

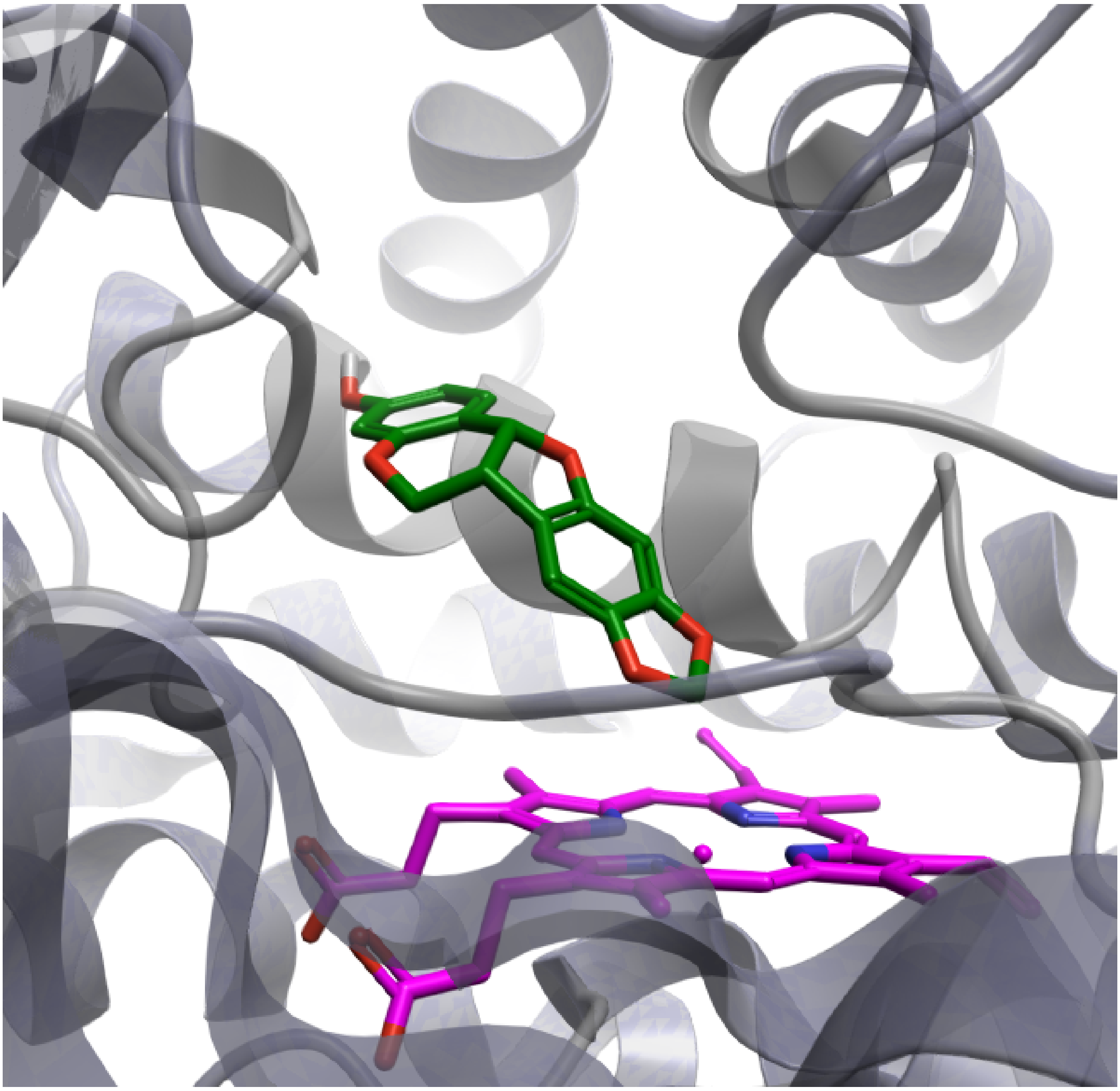

**Figure S7.** Diagram of the probable complex structures formed by CYP124 (structure's PDB ID 6T0G) with Maackiain. The protein is depicted as a cartoon (grey) with the ligand represented as sticks (carbon – green, oxygen – red, hydroxyl's hydrogen – white), with heme shown in stick representation (carbon and ion – magenta, oxygen – red, nitrogen – blue).

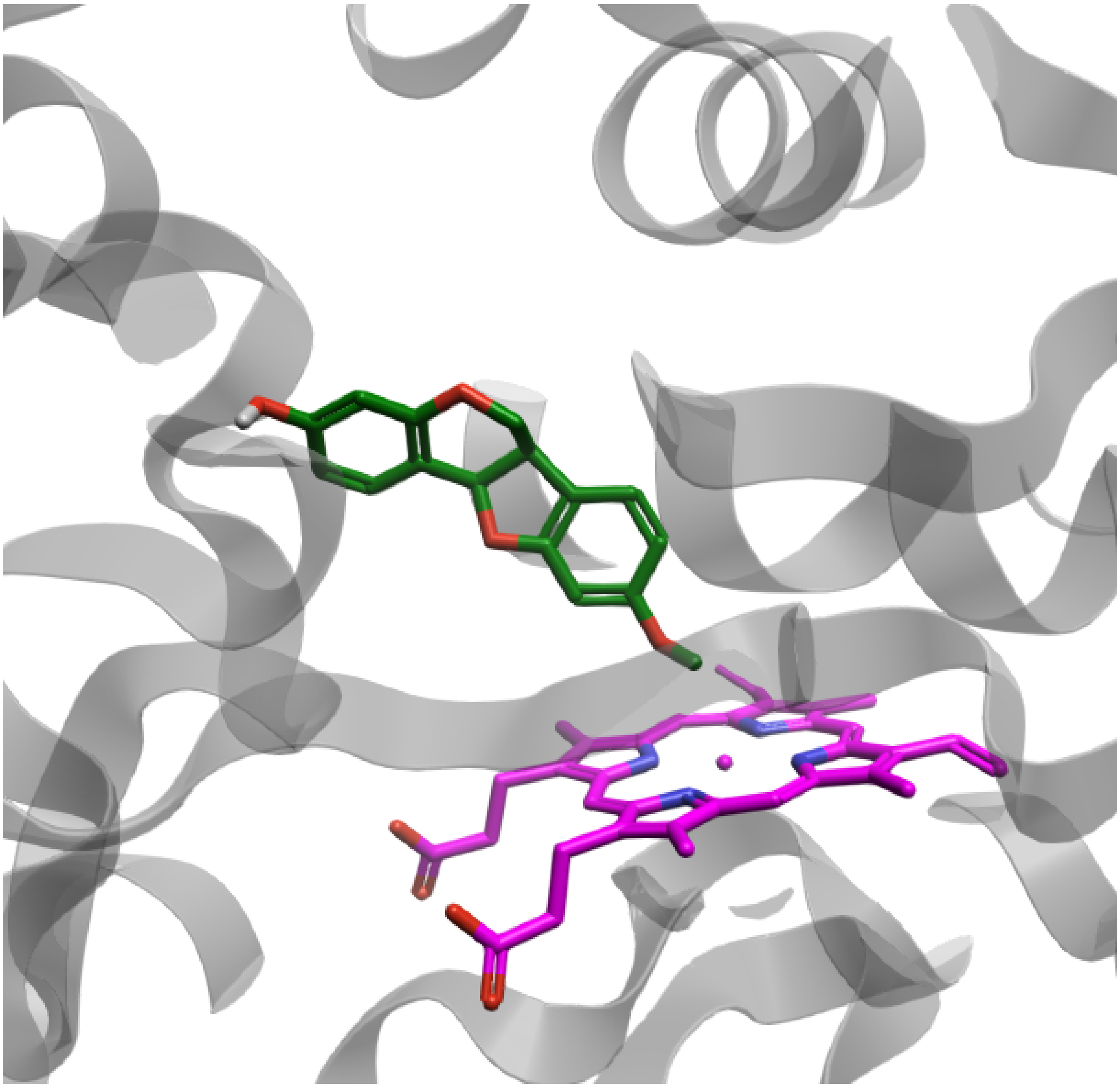

**Figure S8.** Diagram of the probable complex structures formed by CYP124 (structure's PDB ID 6T0K) with Medicarpin. The protein is depicted as a cartoon (grey) with the ligand represented as sticks (carbon – green, oxygen – red, hydroxyl's hydrogen – white), with heme shown in stick representation (carbon and ion – magenta, oxygen – red, nitrogen – blue).

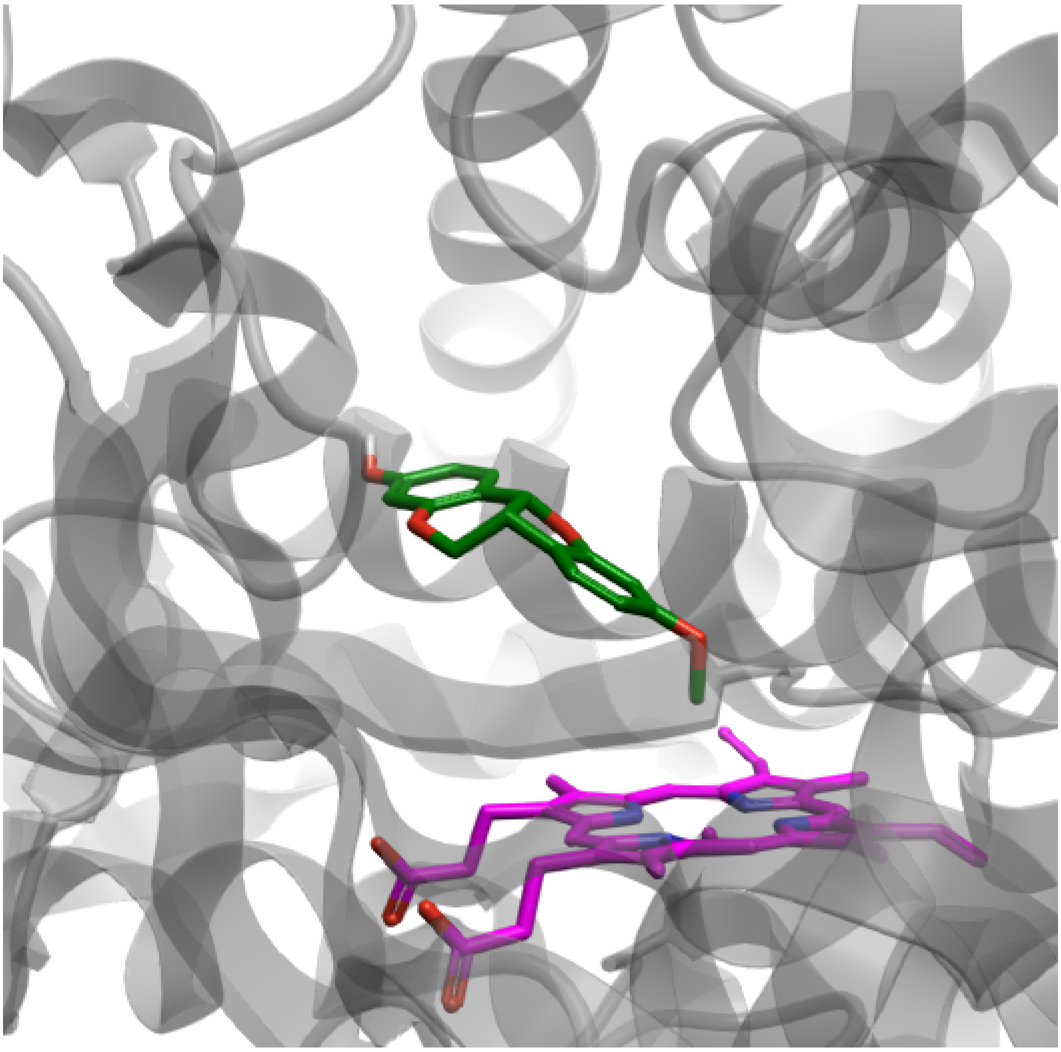

**Figure S9.** Diagram of the probable complex structures formed by CYP124 (structure's PDB ID 6T0F) with Medicarpin. The protein is depicted as a cartoon (grey) with the ligand represented as sticks (carbon – green, oxygen – red, hydroxyl's hydrogen – white), with heme shown in stick representation (carbon and ion – magenta, oxygen – red, nitrogen – blue).

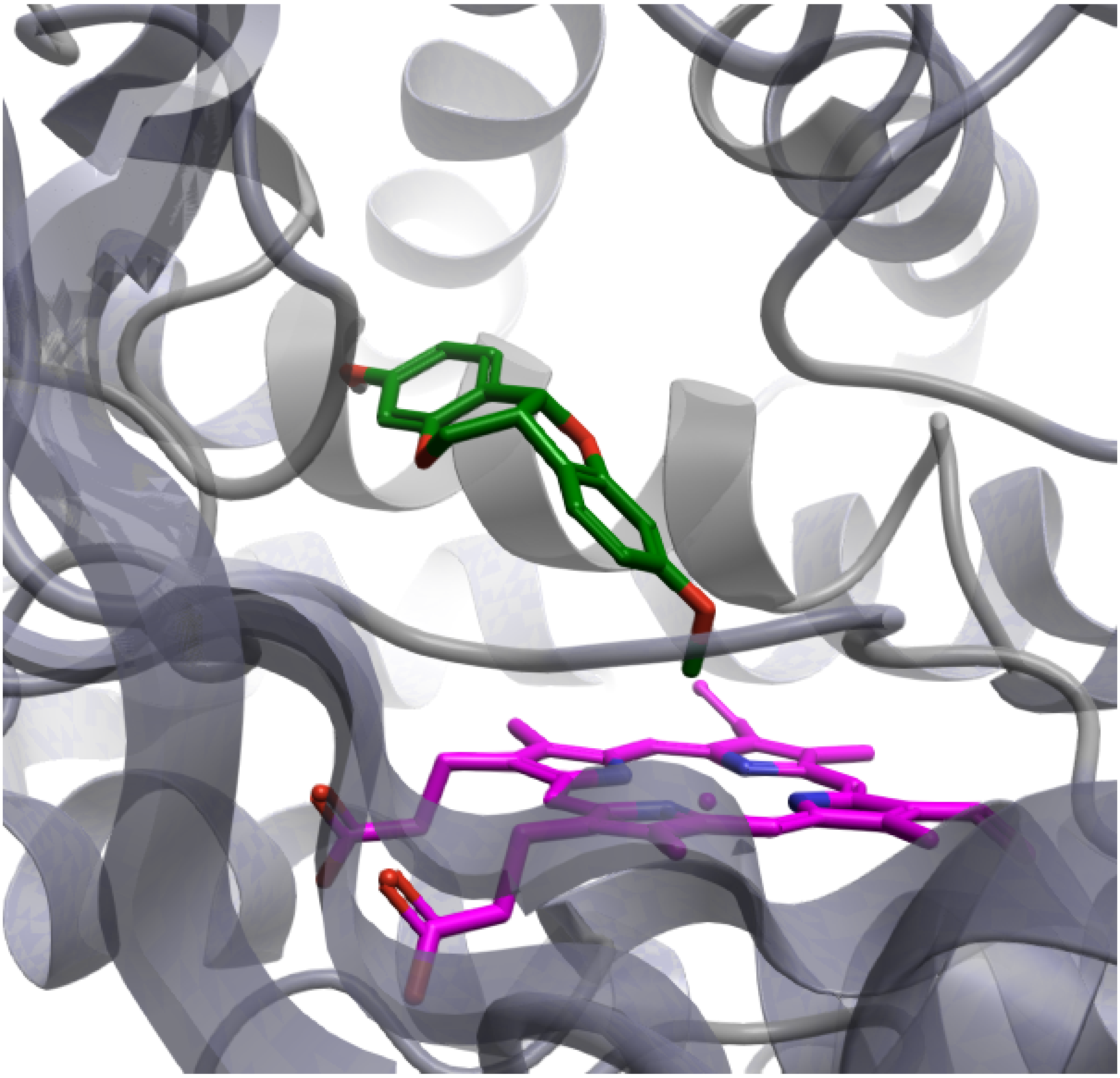

**Figure S10.** Diagram of the probable complex structures formed by CYP124 (structure's PDB ID 6T0G) with Medicarpin. The protein is depicted as a cartoon (grey) with the ligand represented as sticks (carbon – green, oxygen – red, hydroxyl's hydrogen – white), with heme shown in stick representation (carbon and ion – magenta, oxygen – red, nitrogen – blue).

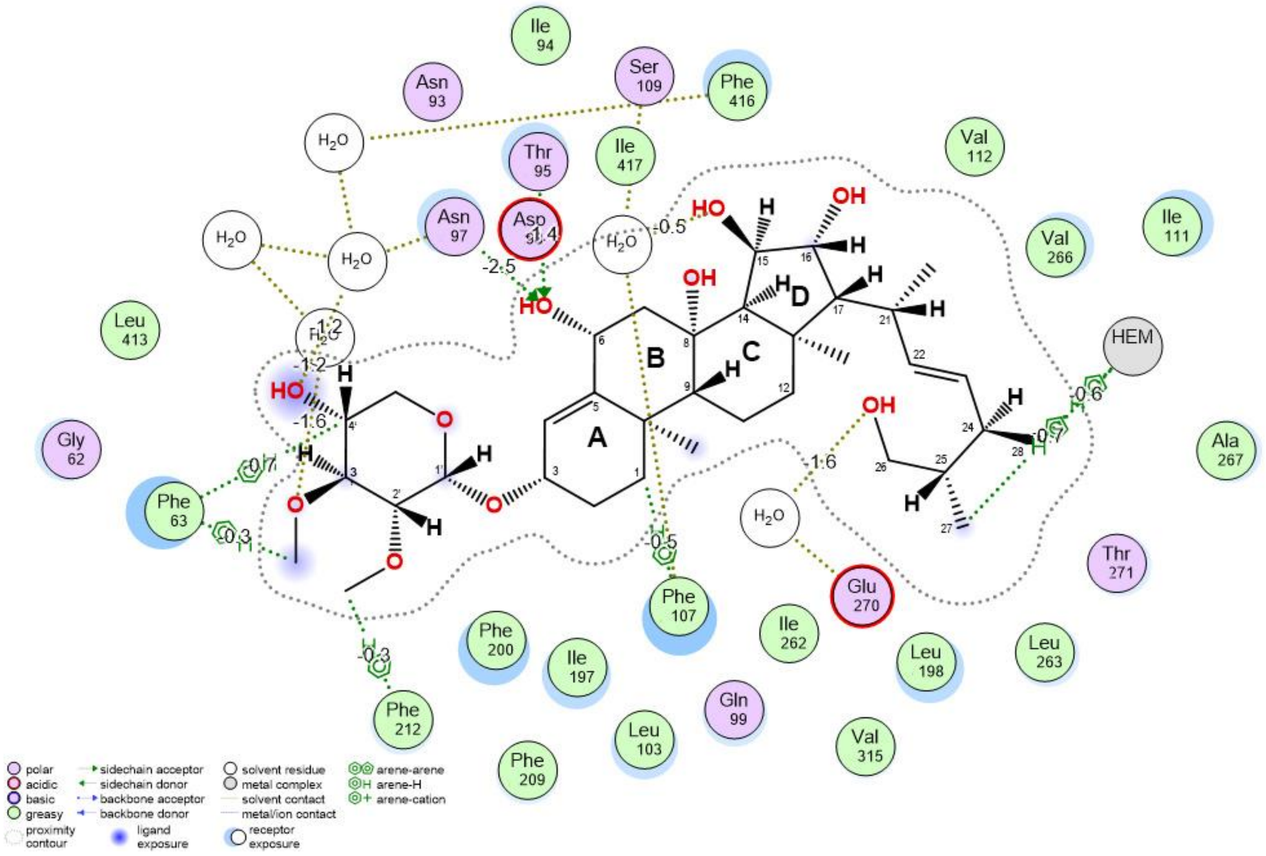


**Figure S11**. 2D diagram of non-covalent interactions of HD-4 with CYP124 of *Mycobacterium tuberculosis* H37Rv. The direct hydrogen bonds are depicted as green dashed lines, the brown dashes represent water-mediated hydrogen bonds and the green dashes represent H-pi conjugates. The estimated interaction contributions to ligand biding energy (kcal/mol) are marked as value above the line indicating an interaction (for example -2.5).

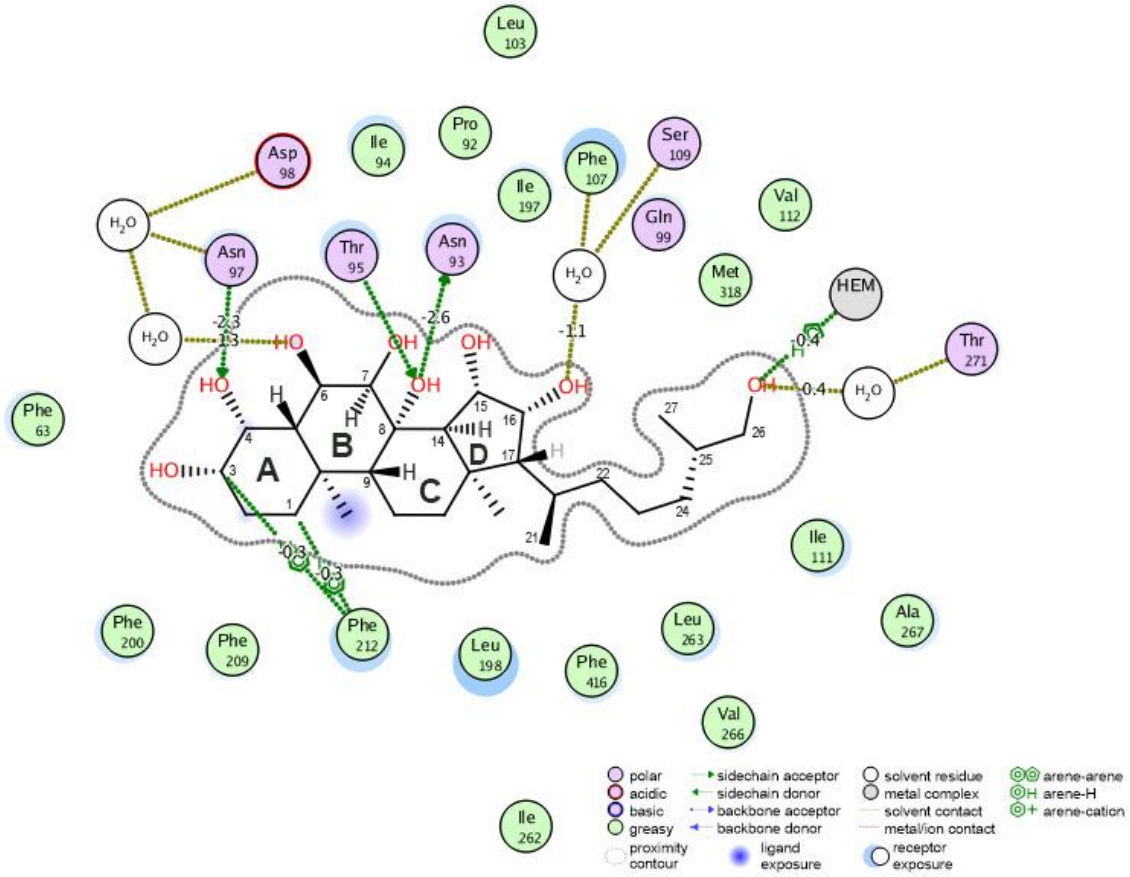


**Figure S12**. 2D diagrams of non-covalent interactions of 15β-octaol with CYP124 of *Mycobacterium tuberculosis* H37Rv. The direct hydrogen bonds are depicted as green dashed lines, the brown dashes represent water-mediated hydrogen bonds and the green dashes represent H-pi conjugates. The estimated interaction contributions to ligand biding energy (kcal/mol) are marked as value above the line indicating an interaction (for example -2.5).

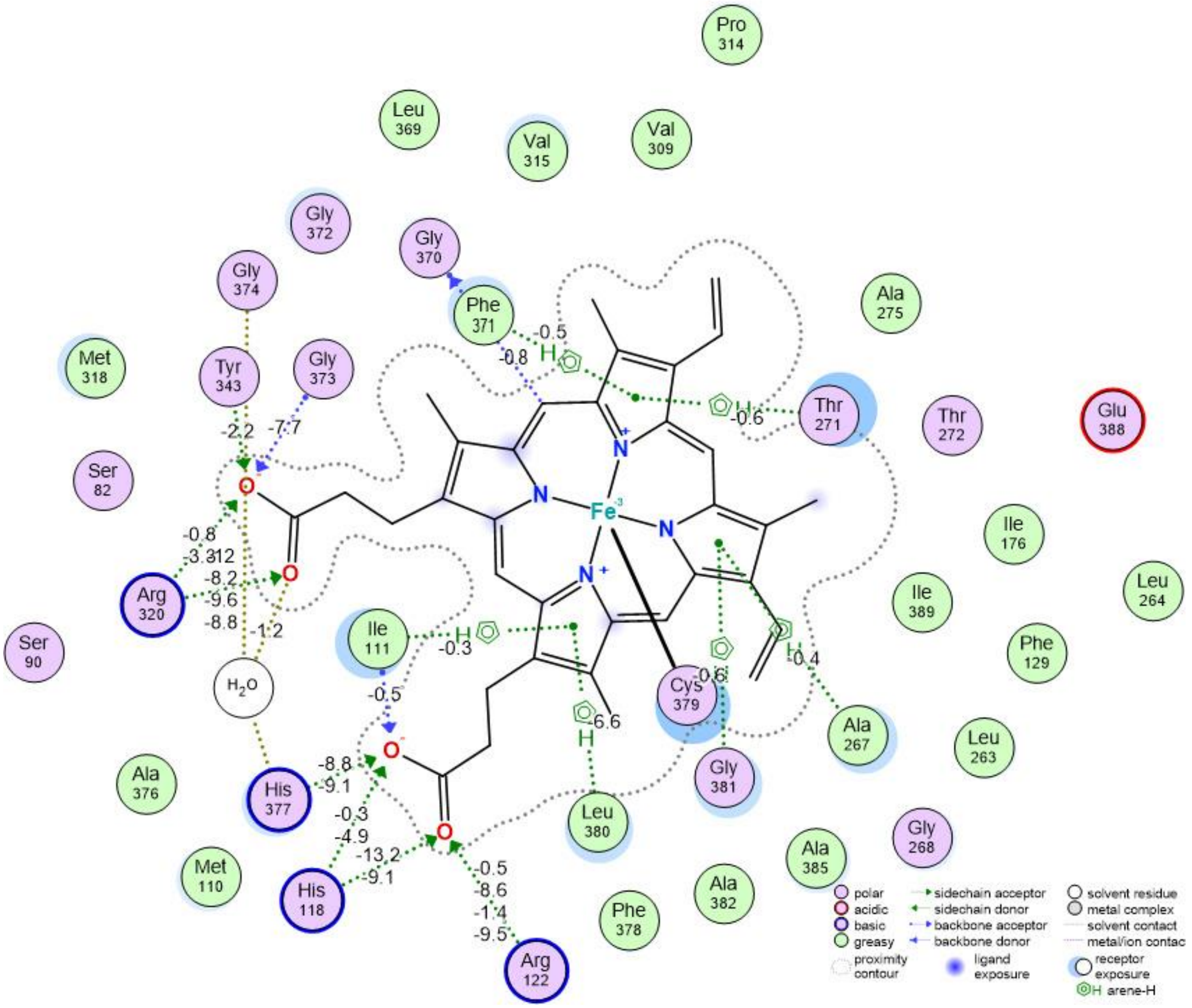


**Figure S13**. 2D diagram of the heme interaction patterns with CYP124 in HD-4 complex. The estimated interaction contributions to ligand biding energy (kcal/mol) are marked as value above the line indicating an interaction (for example -2.5).

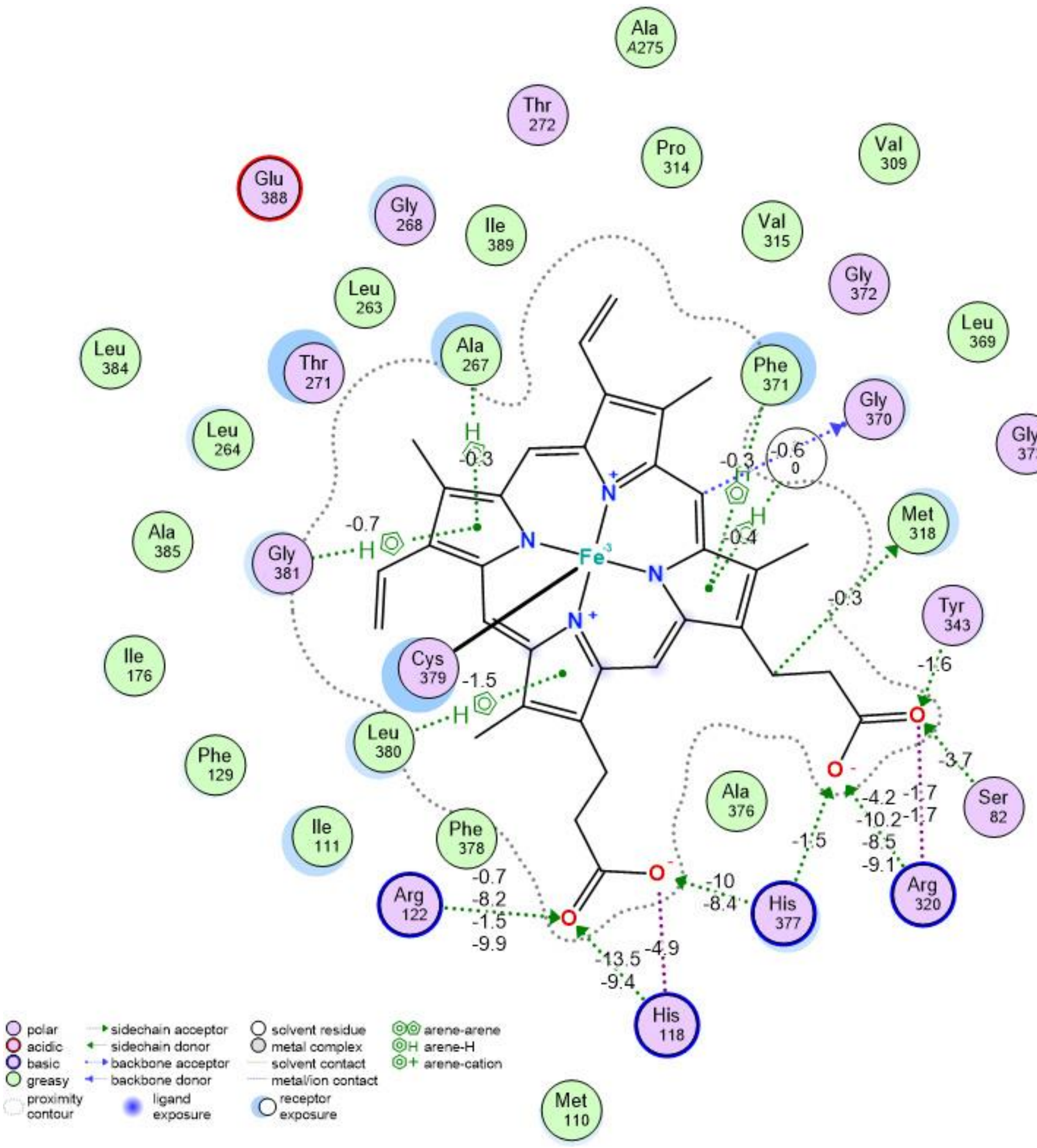


**Figure S14.**. 2D diagrams of the heme interaction patterns with CYP124 in 15β-octaol complex. The estimated interaction contributions to ligand biding energy (kcal/mol) are marked as value above the line indicating an interaction (for example -2.5).
